# DySCo: a general framework for dynamic Functional Connectivity

**DOI:** 10.1101/2024.06.12.598743

**Authors:** Giuseppe de Alteriis, Oliver Sherwood, Alessandro Ciaramella, Robert Leech, Joana Cabral, Federico E. Turkheimer, Paul Expert

## Abstract

A crucial challenge in neuroscience involves characterising brain dynamics from high-dimensional brain recordings. Dynamic Functional Connectivity (dFC) is an analysis paradigm that aims to address this challenge. dFC consists of a time-varying matrix (dFC matrix) expressing how pairwise interactions across brain areas change with time. However, the main dFC approaches have been developed and applied mostly empirically, lacking a unifying theoretical framework, a general interpretation, and a common set of measures to quantify the dFC matrices properties. Moreover, the dFC field has been lacking ad-hoc algorithms to compute and process the matrices efficiently. This has prevented the field to show its full potential with high-dimensional datasets and/or real time applications.

With this paper, we introduce the Dynamic Symmetric Connectivity Matrix analysis framework (DySCo), with its associated repository. DySCo is a unifying approach that allows the study of brain signals at different spatio-temporal scales, down to voxel level, that is computationally ultrafast. DySCo unifies in a single theoretical framework the most employed dFC matrices, which share a common mathematical structure. Doing so it allows:

1) A new interpretation of dFC that further justifies its use to capture the spatiotemporal patterns of data interactions in a form that is easily translatable across different imaging modalities.
2) The introduction of the the Recurrence Matrix EVD to compute and store the eigenvectors and eigenvalues of all types of dFC matrices in an efficent manner that is orders of magnitude faster than naïve algorithms, and without loss of information.
3) To simply define quantities of interest for the dynamic analyses such as: the amount of connectivity (norm of a matrix) the similarity between matrices, their informational complexity.

The methodology developed here is validated on both a synthetic dataset and a rest/N-back task experimental paradigm – the fMRI Human Connectome Project dataset. We demonstrate that all the measures proposed are highly sensitive to changes in brain configurations. To illustrate the computational efficiency of the DySCo toolbox, we perform the analysis at the voxel-level, a computationally very demanding task which is easily afforded by the RMEVD algorithm.

## 2 Introduction

The brain is increasingly recognized as a complex system whose activity is characterized by transient patterns of interaction across its segregated regions ([1]). The field of dynamic Functional Connectivity (dFC) aims to investigate the dynamics of these interactions ([2]). There is a plethora of publications employing methods to uncover the structure of dFC, especially in fMRI ([3, 4, 5]). Starting from a seminal work on schizophrenia ([6]), the field of “Chronnectomics” has shown great potential in elucidating and characterizing differences in typical versus atypical brain dynamics ([7]). Some exemplar applications are in psychiatric disorders ([8, 9, 10, 11]), neurodevelopment ([12]), ageing and neurodegeneration ([13, 14, 15, 16]), cognitive performance and flexibility ([17]), the effect of psychedelics ([18]), and neurological conditions, like epilepsy ([19]) or traumatic brain injury ([20]).

There is also a growing corpus of dFC applications in electrophysiology ([21] extensively reviews its associated methods) and in combined EEG-fMRI data ([22, 23] investigate the link between eeg and bold dFC). In wide-field calcium imaging, dFC has has been shown effective in encoding spontaneous behaviour ([24]) and predicting learning rates in mice (de Alteriis et al. in preparation). These works suggest that brain dynamics and their dFC encoding translate both across species and imaging modalities. This is rendered possible as dFC aims at capturing the dynamic changes in brain-wide global configurations, in contrast to methods that focus on activity in individual regions as mapping the specific role of each region appears to be less important than understanding how they interact.

The foundation of all dFC approaches is a time-varying matrix which expresses how pairwise interactions between nodes of the network change with time in a given brain recording. This significantly improves the classic view of Static Functional Connectivity in which a unique, averaged, connectivity matrix is representative of the whole recording ([25]).

The most common approach to compute dFC is based on sliding window correlation/covariance matrices ([2, 4, 26, 27, 28]). This approach is simple to implement, interpret and can be generalised to any type of signal, from fMRI to electrophysiology, while not over-imposing any hypothesis/model on the signals themselves.

More recently, a method based on instantaneous co-fluctuation patterns has been introduced ([29, 30, 31]). This method can be seen as an instantaneous covariance matrix. These co-fluctuation patterns also hold potential for expanding the investigation into higher-order interactions ([30, 32]). Another approach, which is complementary to these correlative approaches, models brain areas as oscillators with an amplitude and a phase, and computes a measure of phase coherence ([33, 17]). This switches the problem from one of identification of an appropriate time-window size to one of finding a bandwidth of interest, where signals can be legitimately approximated by simple oscillations ([34]). This approach has been especially successful in fMRI signals, which are usually narrow-band, and its application to the study of wake-sleep ([35]), psychedelics ([36, 18]), neurodevelopment ([12]), psychiatric disorders ([11]), ageing ([15]).

These three approaches have methodological commonalities, and a set of operations on the dFC matrices are performed. Firstly, the eigenvectors of dFC matrix are commonly calculated ([17]), either as a dimensionality reduction or as a denoising step. Secondly, a measure that synthesises the overall amount of interactions expressed by the matrix is needed (for example, mean synchrony [37]). Third, a measure of similarity/distance between two dFC matrices is needed either to perform clustering ([4]) or to analyze how dFC changes through the recordings ([13]). Finally, given the increasing attention towards measures of brain complexity, different measures of entropy have been proposed ([38]). All these operations are computationally demanding: a recording of *N* signals at *T* time points produces *T* matrices of size *N* × *N*, which could be hard to process and interpret when spatial and temporal resolution are high. This is why dFC analyses are often carried out on parcellated data.

The results produced in the field of dFC in the past years are mostly empirical, and a formal mathematical framework underpinning the link across the different types of matrices outlined above, and the meaning of their eigenvectors, is lacking. We also believe there is a need to complement the dominant view of dFC, which focuses on tracking changes in functional connections over time, with the identification and characterisation of spatiotemporal statistical patterns of signals. We thus posit that measures of overall connectivity, similarity, and entropy, should be included in the dFC analysis framework. Finally, the dFC analysis framework should explicitly be designed to optimize computational times, allowing real-time computations and parcellation-free analyses, which are prohibitive with current algorithms. This last aspect is critical given the explosion of temporal and spatial scales of neural databases of the latest years, for example, ultrafast fMRI ([39]), high-resolution electrophysiology ([40]), widefield calcium imaging ([24]), 2-photon imaging ([41]).

We propose here the DySCo analysis framework (Dynamic Symmetric Connectivity Matrix Analysis framework). DySCo proposes a unified mathematical formalism and a set of measures and algorithms to compute and analyse dFC matrices. In a nutshell, this comprises: a unique algorithm (the Recurrence Matrix EVD – RMEVD) to compute and store the dFC matrices with their eigenvectors and eigenvalues, that outperforms existing methods in computational speed and memory requirement by orders of magnitude.; and a common set of measures to quantify the evolution of dFC in time. These measures allow to perform the analyses that are typically performed in dFC studies, as described above. DySCo is applicable to any source of data structured as a multivariate time-series. DySCo works seemlessly with very high-dimensional data, both in time and space, and thus paves the road to efficient and systematic as well as the real-time exploration of dynamic patterns of connectivity.

DySCo can also be seen as a (linear) toolbox to do “statistical mechanics” of the brain signals. Correlation/covariance matrices are capturing the shape of the a cloud of data. Their eigenvectors are the principal axes of variation, and the eigenvalues are their importance in describing the cloud of data, in analogy to PCA. With DySCo, we are measuring how this cloud of points evolves in time, i.e., how the spatiotemporal structure, or joint statistics, of the data changes over time. We believe that the statistical mechanics analogy is particularly compelling because in statistical mechanics, the element of interest is not the individual trajectory of a single particle, but the statistical properties of the particles traces and their interactions. The spectral properties of an FC matrix is a convenient and compact representation of this information, i.e. the shape of the data. It is not a case that FC matrices have a similar form to the density matrix in quantum thermodynamics. At at a practical level this analogy may be useful in all the cases where the focal point is the properties of a brain state, rather then the individual trajectories, for example, wake/sleep alternation ([35]), drugs ([36], psychiatric conditions ([42]), ageing ([15]).

## 3 Theory

### 3.1 DySCo

From now on we will refer to any matrix representing dynamical Functional Connectivity (dFC matrix) as **C**(*t*).

DySCo proposes a unified framework to compute **C**(*t*) matrices, their eigenvectors, and metrics of interest: the norm of the **C**(*t*) matrix (see 3.6.1), which is related to the amount of instantaneous interactions, the distance between **C**(*t*) matrices (3.6.3), to perform clustering or to analyze how spatiotemporal patterns change in time, and the **C**(*t*) entropy, which is related to how multidimensionally the signals explore their space (3.6.4). The DySCo framework is based on the Recurrence Matrix EVD (RMEVD), a new algorithm which is order of magnitudes faster than current eigendecomposition methods. DySCo unifies the treatment the different dFC matrices: co-fluctuation, phase locking, correlation, covariance matrices ([15]).

We developed the code to compute all the DySCo quantitities both in MAT-LAB and in Python, which is available at *https://github.com/Mimbero/DySCo*.

### 3.2 Mathematical structure of the DySCo dFC matrices

We will show that all **C**(*t*) matrices have the same structure and can therefore be treated within the same mathematical framework. Given a multivariate timeseries of dimension *N*, all above matrices can be expressed as a dyadic sum, i.e.

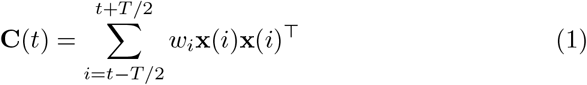

where the vectors **x**(*i*) are a representation of the signals at a time *i*, see the Appendix for the full derivation. Given this structure, **C**(*t*) matrices are:

- symmetric
- positive-semidefinite
- low-rank: the rank is not larger than *T*
- of fixed trace, which is equal to the number of signals *N*. This property holds only for a subset of **C**(*t*) matrices: the sliding window correlation matrix, the “tapered” window correlation matrix, the “instantaneous Phase Alignment” matrix, see below.

We now explicitly write the dFC matrices **C**(*t*) in the format of Equation 1:

- **Sliding window Correlation Matrix**: In this approach, correlations are computed in a window of size *T*:

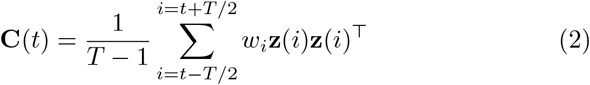

Where **z**(*i*) are the z-scored signals in the window [*t* − *T/*2, *t* + *T/*2]. This is the most commonly employed **C**(*t*) matrix ([24, 43], de Alteriis et al, in preparation) together with the sliding window covariance matrix ([4, 44]). Windows can be “square” if *w*_*i*_ = 1 ∀ *i* or “tapered”/”weighted” if the weights *w* form a window with smooth edges ([2]).

- **Sliding window Covariance Matrix**: Similarly to the correlation case, the covariances are computed in a window of size *T* with weights *w*_*i*_, **y**(*i*) are the demeaned signals in the window.

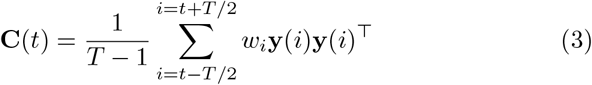

- **Cofluctuation Matrix**: The co-fluctuation matrix can be conceived as a sliding window covariance matrix with window-size *T* = 1:

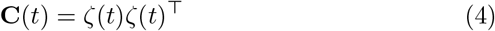

where ζ is the z-scored signal in the whole recording. Co-fluctuation matrices have the property that the static connectivity matrix is the average of all the instantaneous co-fluctuation matrices ([29]).

- **Instantaneous Phase Alignment Matrix**: The instantaneous Phase Alignment Matrix (**iPA**)([17, 42, 15]) quantifies the extent to which signals are in phase. The Phase Alignment Matrix assumes that every brain area can be modeled as an oscillator with an instantaneous phase *θ*(*t*). This is true for narrow-band signals, like fMRI, or signals filtered in a specific band (for example EEG filtered in different bands). The extraction of the instantaneous phase requires the use of the Hilbert Transform ([15, 33]) or Wavelet Transform ([45]). The “instantaneous Phase Alignment” measures the cosine of the phase difference of the signals. The vectors **c**(*t*) and **s**(*t*) are respectively the element-wise cosine and sine of the instantaneous angles ([18, 15]). It is worth noting that the **iPA**(*t*) matrix is analogous to the correlation matrix under the local approximation of signals with sinusoids, see Appendix. Therefore, the **iPA**(*t*) matrix has the same role as the sliding window correlation matrix. However, the sliding window computes correlations in a window, while the **iPA**(*t*) matrix assumes that the signals are locally approximated by simple oscillations.

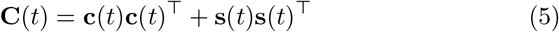

### 3.3 Interpretations of dFC

Dynamic Functional Connectivity is usually conceived as study of the changes in the connection strength between nodes in a brain functional network (irrespective of whether or not there is an anatomical circuitry underneath) ([2, 46]).

We add here a complementary interpretation of dFC. Equation 1 shows that **C**(*t*) is a statistical representation of spatial pattern at a time *t* estimated over a time window of size *T* (in case of **iPA** and cofluctuation, *T* = 1). The evolution of *C*(*t*) therefore represents the evolution of the spatio-temporal patterns. The size of the window determines the trade-off between the accuracy of the estimation of the statistical pattern and the resolution in time. In general, the choice of the window depends on the properties of the signals and cannot be determined a-priori.

This interpretation has the following implications:

- **C**(*t*) is a way to approximate the dynamics of a complex system in a datadriven way, without over-imposing any model or theoretical bias on the processing of the signals. Indeed, the only assumption is that in every window the signals are sufficiently stationary to be well described by their second order statistics. This is generally not true in the whole recording, as for example task and rest experimental blocks would have different characterisation, see Fig. 1.B.ii. Thus, **C**(*t*) expresses how these statistics change with time. Most complex systems move in a landscape of different spatiotemporal interaction patterns ([47]). **C**(*t*) is able to capture this movement (see Fig. 1 for a toy example).
- Since **C**(*t*) can be seen as an average of pairwise products, as suggested by [24], we can interpret **C**(*t*) as the second-order Taylor expansion of any function of brain data. Indeed, we can formally write any brain output, for example behaviour, as *b*(*t*) = *f* (**x**(*t*)), where **x**(*t*) is brain activity and *f* a function mapping activity to behaviour. Using a second-order Taylor expansion in *t*_0_, we can write 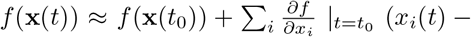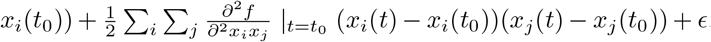 Overall, this suggests that **C**(*t*) can be seen as the quadratic term of a second order approximation of brain function (see [24] for the full development).
- As all **C**(*t*) are symmetric matrices, it is natural characterise them with their eignedecomposition. And, since they are low-rank matrices, a small number, Rk(**C**(*t*)) ≤ *T* ≪ *N*, of eigenvectors suffice to express all the information contained in the matrix:

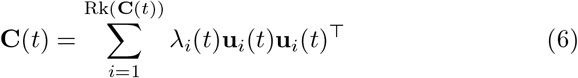
- One could argue that for a too small window, **C**(*t*) is not able to capture neither true correlations/covariances/phase locking patterns, nor the underlying network structure, but the eigenvectors, respectively of the size of the window, are the principal axes of variation in the data. Even in a 1 time frame window, the main axis of variation of the signal is the signal itself. Finally, the eigenvector representation of **C**(*t*) allows a straightforward strategy for denoising the signal, by retaining a subset of eigenvectors associated to the largest eigenvalues, which are proportional to the amount of variance expressed.

**Figure 1:**
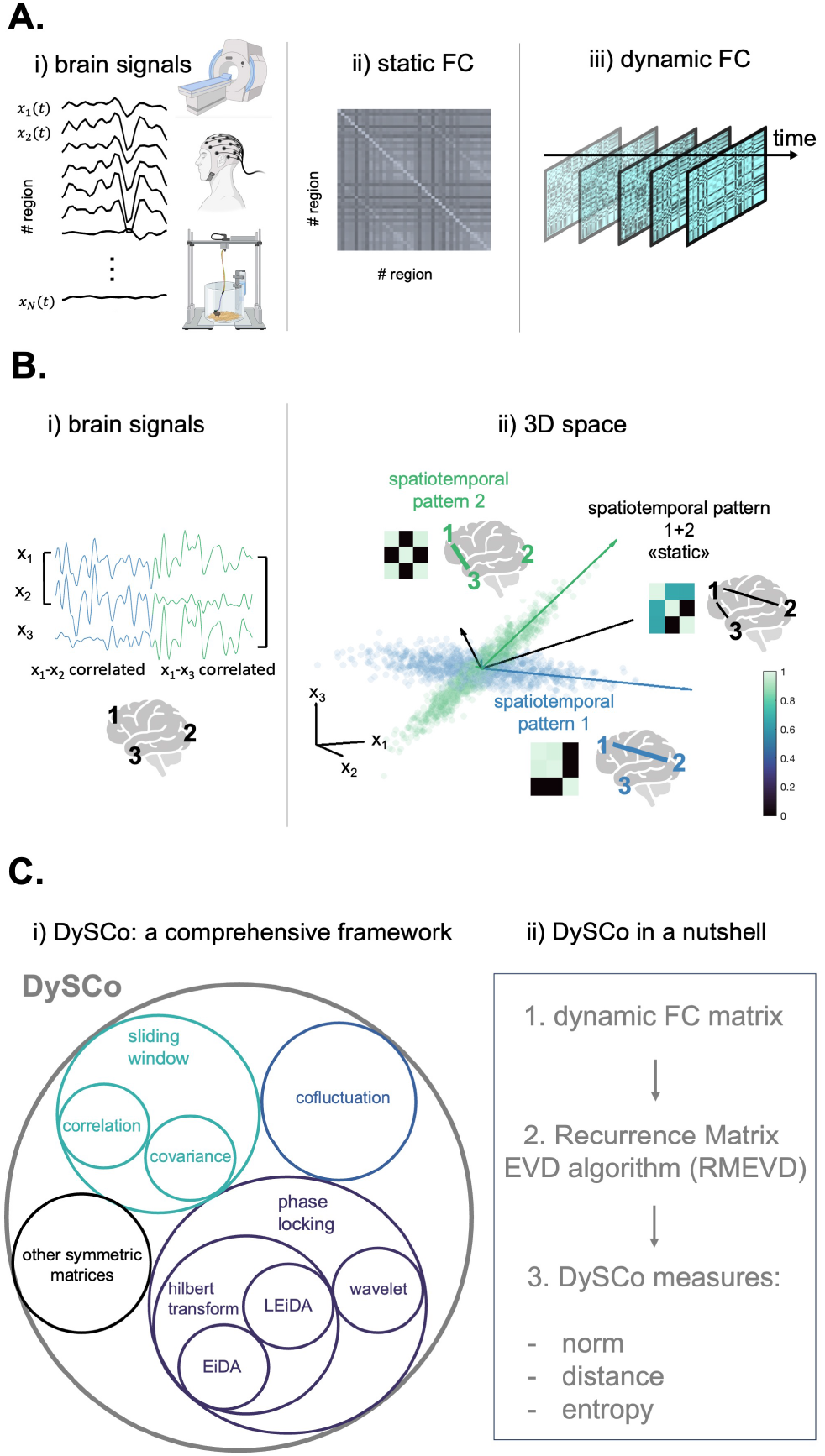
**A**. What is dynamic Functional Connectivity: i) We can start from any set of brain recordings, where each signal is referred to a brain location (e.g. fMRI, EEG, intracranial recordings in rodents, and more). ii) it is well known that “static” Functional Connectivity (FC) is a matrix where each entry is a functional measure of interaction between two regions, for example, the Pearson Correlation Coefficient. iii) Dynamic Functional Connectivity (dFC) is a FC matrix (that can be calculated in different ways, see below) that changes with time, under the assumption that patterns of brain interactions are non-stationary. **B**. Why dFC is important: i) In this toy example, 3 brain signals are recorded, referred to 3 anatomical locations (*x*_1_(*t*), *x*_2_(*t*), *x*_3_(*t*)). In the first half of the recording (blue half) *x*_1_(*t*) and *x*_2_(*t*) are highly correlated (high FC), while in the second half *x*_1_(*t*) and *x*_3_(*t*) are highly correlated. Thus, the brain switches between two different spatiotemporal patterns of interaction (pattern 1, blue, and pattern 2, green). ii) Pattern 1 can be seen as a matrix, as a graph, and as a set of main axes of variation in a 3D space (blue), and the same for pattern 2 (green). In this toy example, the switch from a high 1-2 correlation to a high 1-3 correlation can be seen as a change in the connectivity matrix or a rotation of the main axes of variation of the signals in the 3D space. However, by using a “static” approach, this switch would not be captured, and a spurious spatiotemporal pattern (the black one), associated with a spurious set of axes of variation, would appear, which does not reflect any actual brain configuration. This is why dFC is a tool to investigate brain dynamics, by looking at how spatiotemporal patterns of interaction change in time. **C**. The Dynamic Symmetric Connectivity (DySCo) Matrix analysis framework for dFC: i) DySCo is a comprehensive framework that puts together different dFC approaches. Interestingly, these include the 3 most employed methods for dFC, i.e., sliding window correlation/covariance, co-fluctuation, approaches based on phase locking. They all involve symmetric matrices. ii) DySCo proposes a unified mathematical formalism and a set of measures and algorithms to compute and analyse dFC matrices. In a nutshell, this entails: 1. The selection of a dFC matrix 2. A unique algorithm (the Recurrence Matrix EVD) to compute and store the dFC matrices with their eigenvectors and eigenvalues, which is orders of magnitude faster and more memory efficient than naïve approaches 3. A common set of measures to quantify the evolution of dFC in time. These measures allow to perform the analyses that are typically performed in dFC studies. They can be classified in three categories: measures based on the total amount of dynamic interactions (matrix norm); measures based on distance/similarity of dFC patterns (e.g. to perform clustering); measures based on the entropy of the dFC patterns.

### 3.4 On the importance of the Leading Eigenvector

The Leading Eigenvector is associated to the largest eigenvalue of **C**(*t*) and is widely employed in both phase-locking studies ([17, 12, 18]) and correlation studies (in this case also known as Eigenvector Centrality –[48]). Its importance can be summarised as it being the best single vector approximation of a dFC matrix:

- It is the main axis of variation of the signals in the window/bandwidth of observation (dominant mode of dynamic connectivity). Each of its entries refers to an anatomical location, so the leading eigenvector is useful to visualize the dominant mode in the anatomical space ([49]).
- The outer product of the leading eigenvector with itself approximates **C**(*t*). Indeed, if 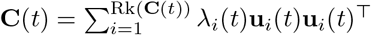, then **C**(*t*) ≈ *λ*_1_(*t*)**u**_1_(*t*)**u**_1_(*t*)^⊤^. This is also used to measure “eigenvector centrality”, because it is a measure of how “central” an area is (i.e. how much it engages interactions with the others). In the anatomical space, an intuitive way to read the leading eigenvector is the following: areas with the same sign have a positive interaction (i.e. positive correlation, covariance, etc), while areas with opposite signs have a negative interactions. The amplitude of the leading eigenvector indicates the centrality of the area.
- Following the same logic, the leading eigenvector of the **iPA** matrix can be seen as the main mode of oscillation, since the **iPA** matrix expresses co-oscillations rather than correlations. Multiple studies have shown that the brain explores a set of oscillatory modes, which can be computed by performing temporal clustering (e.g. k-means) of the leading eigenvectors of the **iPA** matrix. This is known as the LEiDA method (Leading Eigenvector Dynamics Analysis)([17, 37]).

### 3.5 The Recurrence Matrix EVD algorithm for the computation of dFC matrices and their eigenvectors

In this subsection, we introduce the methodological core of the DySCo framework, that allows for ultra fast eigendecompostion of very high dimensional matrices.

Let us consider a generic **C**(*t*) matrix that is expressed as a dyadic sum 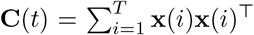. Being a matrix, **C**(*t*) is a linear operator from ℝ^*N*^ to ℝ^*N*^, where *N* is the number of signals. The fact that a dFC matrix is a sum means that all its outputs lie in the space spanned by the *T* vectors **x**(*i*). This implies that its rank Rk is not higher than *T*, so Rk ≤ *T*. Indeed, the rank of a linear operator is the dimensionality of the space where it maps its inputs. Moreover, this implies that the eigenvectors of a dFC matrix must be a linear combination of the *T* vectors, and the eigenvectors associated with non null eigenvalues will be no more than the rank, so no more than *T*. This means that any dFC matrix is an extremely low ranked operator, provided *T* ≪ *N* (which is the case in practical applications). Thus, Rk ≤ *T* ≪ *N*, and the full information of the dFC pattern can be stored without approximation, or losslessly, in at most *T* eigenvectors.

Moreover, the Rk eigenvectors and their associated eigenvalues can be computed using the Recurrence Matrix EigenVector Decomposition algorithm (RMEVD) (see Appendix):

- for a generic dFC matrix at a generic time point *t* **C**(*t*) which is expressed as 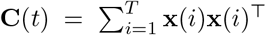 we define the Recurrence Matrix **R**(*t*)_*ij*_ = **x**(*i*)^⊤^**x**(*j*). This matrix of scalar products has the size *T* × *T*
- The eigenvalues of the **C**(*t*) matrix are the eigenvalues of the Recurrence Matrix **R**(*t*).
- Each eigenvector **v** of the Recurrence Matrix is a *T* –tuple of coefficients. The eigenvectors of the **C**(*t*) matrix are a linear combination of the **x** vectors, where the coefficients are **v**_*i*_.

Note that the **iPA**(*t*) matrix, (see Eq. 5), is rank 2 and therefore its associated Recurrence Matrix is a 2 × 2 matrix and its eigenvalues and eigenvectors can be computed analytically. For the analytical calculations, and an extended discussion on the **iPA** matrix structure, see [15]. Note also that the sliding window correlation (see Eq. 2) and covariance (Eq. 3) matrices are rank *T* − 1 and not *T*, given that before application of the Equation 1 signals are demeaned and therefore lose a degree of freedom. Finally, note that the co-fluctuation matrix (Eq. 4) is rank 1, as it is just ζ(*t*)ζ(*t*)^⊤^, so its eigenvector is trivially ζ(*t*), and its associated non null eigenvalue 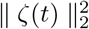

This algorithm implies that the full information of a dFC matrix can be extracted without the need of computing the matrix itself. Its eigendecomposition (EVD) operates on the *T* × *T* recurrence matrix, with associated time complexity of 𝒪 (*NT* ^2^ + *T* ^3^) and space complexity of 𝒪 (*T* ^2^) instead of a *N* ×*N* one, that has associated time complexity of 𝒪 (*N* ^3^) and space complexity of 𝒪 (*N* ^2^)([50], [51]). This can make algorithms for computation and storage orders of magnitude more efficient (see Figure 4, in Section 5.1). We refer the reader to the Appendix for an extended proof that considers the general weighted sum 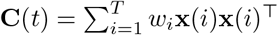

Finally, note that the RMEVD is efficient when *T < N*, which is typically the case in dFC. In case *T > N*, the DySCo framework still applies, however, we suggest using a classic EVD because the RMEVD would not improve the computational speed given that matrices would be full rank.

We observe also that this approach is similar to the Dynamic Mode Decomposition introduced in Fluid Mechanics ([52]).

#### 3.5.1 On the meaning of the Recurrence Matrix

More than being a tool to compute the eigenvectors of the dFC matrix efficiently, the Recurrence Matrix has also a physical meaning for the correlation and covariance matrices: it is the matrix of scalar products, and thus of matching/similarity, of the signals in space, contrary to the correlation matrix that expresses similarity in time. Indeed, within a window, the entry *ij* of the Recurrence Matrix is the scalar product between the signals **x**(*t*_*i*_) and **x**(*t*_*j*_). Physically speaking, the vector **x**(*t*_*i*_) is the collection of all brain signals, at all locations, at time *t*_*i*_, and similarly for **x**(*t*_*j*_). Thus, if the scalar product is high, this means that the brain signals are similar at *t*_*i*_ and *t*_*j*_ – if it is zero, it means that they are orthogonal. The Connectivity Matrix is in the space domain, and quantifies whether a couple of brain areas have a similar time-course. The Recurrence Matrix is in the time domain, and quantifies whether a couple of time points are characterized by a similar whole-brain configuration.

Thus, the eigenvectors **v**, the recurrence modes, of the Recurrence Matrix are time vectors (*i* = 1… *T*) and are associated to the changes in time instead of changes in space of the signals. For example, the dominant recurrence mode, recurrence mode associated with a high eigenvalue, would be a dominant temporal similarity pattern in the signals.

**Table 1:**
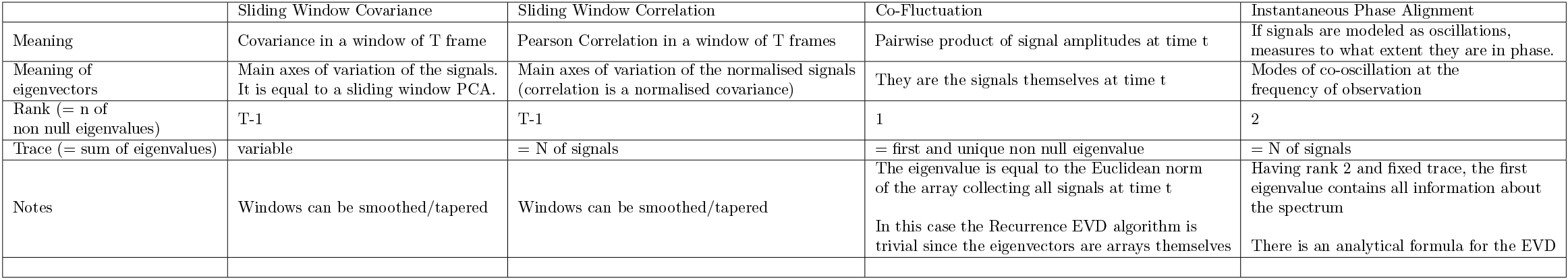
Description of the dFC matrices presented above and their main properties.

### 3.6 DySCo measures

Here, we introduce a set of measure that are aimed at characterising the spatio-temporal patterns captured by the dFC approach and brain activity in general.

#### 3.6.1 Norms

The norm of a **C**(*t*) matrix is a synthetic measure of the overall amount of interactions expressed within ([53]). Here we propose three of the most employed matrix norms in linear algebra. The Recurrence Matrix EVD of the dFC matrices suggested in the DySCo framework allows to compute these norms without computing the matrices directly and storing them.

- The Schatten norm 1 (Trace norm) of a matrix, ∥ **C**(*t*) ∥_1_, is the sum of the absolute values of its eigenvalues: Σ _i_ |*λ*_*i*_(*t*)|
- The Schatten norm 2 (Frobenius norm) of a matrix, ∥ **C**(*t*) ∥_2_, is the square root of the sum of the squared eigenvalues: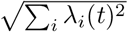 which coincides with the square root of the sum of all the squared values of the matrix 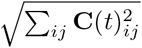 since dFC is symmetric.
- The Schatten norm ∞ (Spectral norm) of a matrix, ∥ **C**(*t*) ∥_∞_, coincides with the largest absolute value of its eigenvalues: max_*i*_ {|*λ*_*i*_(*t*)|}.

Note that, in the specific cases of the correlation matrix and **iPA**, the matrices **C**(*t*) have a fixed trace (see Section 3.2). Consequently, their norm-1 becomes trivial and simply corresponds to their size *N*. Moreover, the cofluctuation matrix has trivially one non-null eigenvalue (see Section 3.5), therefore all the norms coincide, and in that specific case they coincide with the norm of the vector ζ(*t*).

#### 3.6.2 Norm Metastability

Norm Metastability is the standard deviation in time of the norm of the dFC matrix ([15], de Alteriis et al in preparation,[42]).

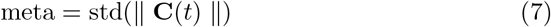

Norm metastability is a measure of variability in the exploration of connectivity patterns. It is a measure of how much the dFC matrix oscillates between time points with high norm and time points with low norm, and reflects thus simultaneous tendencies for coupling and decoupling. In [15], the standard deviation of the infinite norm has been introduced as spectral metastability ([11]).

#### 3.6.3 Distances between dFC operators

If a matrix norm exist, the distance can be computed as just the norm of the difference of two matrices. Therefore:

- distance 1 is ∥ **C**(*t*_1_) − **C**(*t*_2_) ∥_1_
- distance 2 is ∥ **C**(*t*_1_) − **C**(*t*_2_) ∥_2_
- distance ∞ is ∥ **C**(*t*_1_) − **C**(*t*_2_) ∥_∞_

It may also be the case, in some applications, that before computing the distances matrices are normalized to have all unit norm.

Defining a distance between **C**(*t*) matrices is useful for several reasons:

- Clustering dFC patterns is a very common approach in neuroimaging ([4, 17]). To cluster it is necessary to define a distance between **C**(*t*) matrices.
- A distance between dFC patterns allows to compute properties of how they explore the dFC space, like the speed at which the **C**(*t*) pattern evolves with time. The reconfiguration speed for a lag *τ* is defined ([13]):

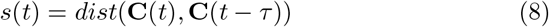
- A distance allows to build the Functional Connectivity Dynamics matrix (FCD). FCD is a time-to-time distance matrix: the entry *ij* of FCD is the distance of **C**(*t*_*i*_) with **C**(*t*_*j*_). This is a condensed potrait of the evolution and properties of dFC in the whole recording ([54]).

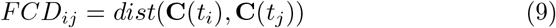

It is still possible to apply the Recurrence Matrix EVD (Section 3.5) to compute the above distances, since they are based on the eigenvalues of the matrix **C**(*t*_1_) − **C**(*t*_2_). It is thus possible to exploit the information expressed in the eigenvectors and compute the distance without rebuilding the matrices. Note that, especially in the case of the norm-2, this turns out to be orders of magnitude faster than the one computed by calculating the full matrix instead of the eigenvector representation, see Appendix.

Note also that, since the correlation and Instantaneous Phase Alignment matrices have all ones on the diagonal, the distance 1 in that case becomes trivially zero – it is therefore suggested in those cases to use the other two distances.

Note: it is a common practice to represent **C**(*t*) matrices as the vectorised **C**(*t*) matrices, or their vectorised upper-triangular part, given the symmetry. One could then think that the matrix distance is equal to the vector distance of the vectorised matrices. However, this is incorrect, because the norm of the vectorised matrix does not coincide with the norm of the matrix, and so does the distance. The only exception is the norm 2, where the two things coincide. However, even in that case, our proposed formulation of the norm 2 is still advantageous because it speeds up computations by orders of magnitude to up to two/three orders of magnitude in practical scenarios.

Another measure commonly employed is the cosine similarity between two vectorized dFC matrices ([13, 54]), which quantifies the matching between two matrices. This is not a distance, but a measure of similarity.

To define a measure of matching it is necessary to define an inner product between two matrices. We introduce the Frobenius Product between two matrices **C**(*t*_1_) and **C**(*t*_2_):

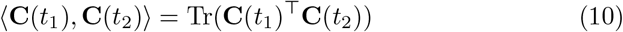

The square root of the Frobenius product of a matrix with itself is its Frobenius norm. So, exactly as in vectors, it is possible to define a measure of alignment:

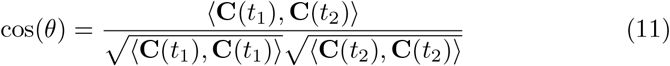

The DySCo framework provides a fast formula also to compute this quantity (see Appendix).

Finally, it may be the case that the quantity of interest is a subset of eigenvectors of the **C**(*t*_1_) and **C**(*t*_2_) matrices, without interest on the eigenvalues. In that case a measure of alignment of the eigenvectors is needed. This measure corresponds to the Frobenius distance between the projector matrices of **C**(*t*_1_) and **C**(*t*_2_) (i.e. the matrices that project, respectively, on the eigenvectors of **C**(*t*_1_) and **C**(*t*_2_)). See Appendix for the full derivation.

We observe that the FCD matrix, being a distance matrix, allows also nonlinear dimensionality reduction techniques like diffusion embedding, as done in ([24]).

#### 3.6.4 Von Neumann entropy of the dFC pattern and its interpretation

The spatiotemporal complexity of brain signals is of great biological interest as it is believed to be related to health and disease as well as consciousness ([55]), sleep-wake transitions ([56, 57]), psychedelics ([58]). We thus include the Von Neumann entropy of **C**(*t*) ([59, 38]):

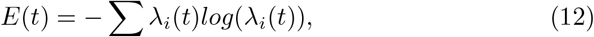

with the eigenvalue spectrum normalized to one.

This measures how broadly the brain is exploring the pattern of its possible configurations (i.e. how multi-dimensionally signals are spanning their axes of variation – eigenvectors) being a measure of variability of the eigenspectrum.

Indeed, the spectrum of eigenvalues associated to the eigenvectors expresses their relative importance. For example, in case where an eigenvector is strongly dominant, the first eigenvalue is high and the others are close to zero. On the other extreme, if all the eigenvectors have the same role, meaning that the brain is exploring its configuration more broadly in the multidimensional space of all its possible configurations, the associated eigenvalues will all have similar magnitudes.

In the first limit case the signal **x**(*t*) in the window would stay always parallel to itself (as a vector), so all the signals are perfectly correlated, all the signals are perfectly oscillating along the axis of variation which is *x* itself. There are no additional axes of variation. On the contrary, the more the spatiotemporal patterns are rich, the more there will be axes of variation, corresponding to more eigenvectors of **C**(*t*) with non-zero eigenvalues. For a visual explanation, see ([15]), where this concept is applied to the **iPA** matrix.

Note that, in case the window is weighted, the eigenvalue spectrum will be forced to make some vectors dominant. The entropy must be considered carefully in this case.

## 4. Materials and Methods

### 4.1 The DySCo repository

The DySCO framework comes with a repository. We developed the code to compute all the DySCo quantitities both in MATLAB and in Python, which is available at *https://github.com/Mimbero/DySCo*. Note that, both in MATLAB and in Python, we provide all the “core functions” (compute RMEVD, compute norm, distance, etc…) to autonomously build a processing pipeline. However, we also offer an already built example Python pipeline, the one that has been used to process the HCP data. The repository also features a Python GUI to run the analyses.

### 4.2 Investigation of computational efficiency of the RMEVD in the DySCo framework

We first investigated if the computational speed-up of the RMEVD algorithm expected from Theory was confirmed. To do so, we fixed a window of length *T* = 10 and we generated *N* Gaussian i.i.d. random signals in the window. We computed the covariance matrix of the signals in the window using both the Recurrence Matrix EVD proposed in DySCo and the standard numerical algorithm provided by MATLAB (using the *eigs* function as in the version 2021A). We computed the time needed to perform EVD in both cases. We varied *N* logarithmically, from *N* = 100 to *N* = 10^4^. We then repeated this procedure 20 times for each *N*. Computation times were calculated on an Apple M2 CPU.

### 4.3 Application to Simulated Data

To validate the method, we generated a synthetic time-series and imposed an underlying time-varying connectivity pattern. More specifically, we generated a 10-dimensional Gaussian random signal with zero mean and unit variance, consisting of 5 chunks, each 1000 time-frames long. We chose Gaussian random signals to be as general as possible. We then generated 5 random covariance matrices, and multiplied each chunk with the Cholesky Decomposition of one of these matrices. This procedure imposed a different covariance matrix on each chunk of the signal.

Following the DySCo framework we computed the sliding window covariance matrix using a window size of 121 frames, (we use odd numbers to make windows symmetric). We computed the reconfiguration speed using the 1,2 and ∞-distances and a *τ* value of 100 frames. Distances were computed on the matrices normalized by their norms. Finally, we computed the Functional Connectivity Dynamics matrices using the distances 1,2 and Infinity.

### 4.4 Application to task fMRI

#### 4.4.1 Summary of the practical steps in the DySCo Framework

It is important to stress that DySCo is not a single method per se, but a unified framework for the dFC methods and measures described in the Theory Section. We illustrate the DySCo pipeline in Figure 3. It consists of three steps:

**Figure 2:**
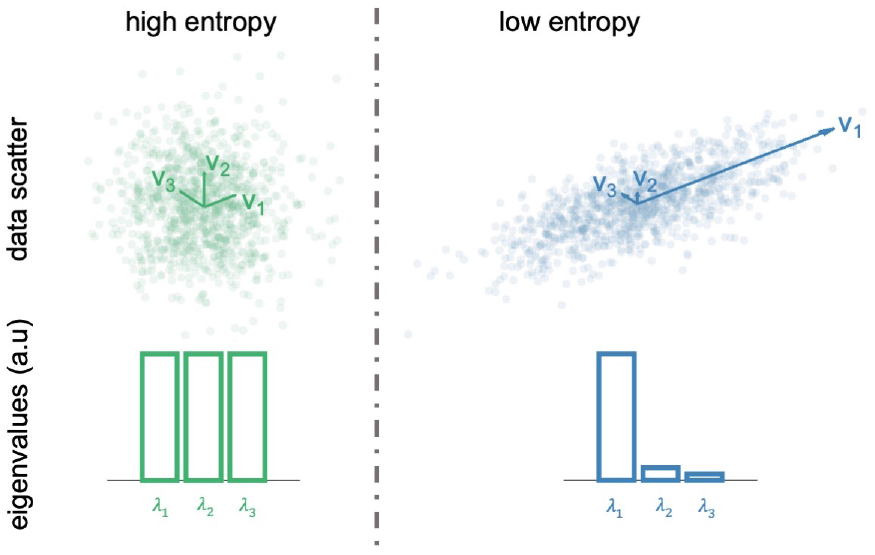
Two example cases to show the Von Neumann Entropy, in an Example 3-dimensional random signal. In the first case, there is no main axis of variation, thus the eigenvalues are all similar. This corresponds to a high Von-Neumann entropy. In the second case, there is a main axis of variation, which corresponds to a less dispersed eigenvalue spectrum. This corresponds to a low Von-Neumann entropy.

**Figure 3:**
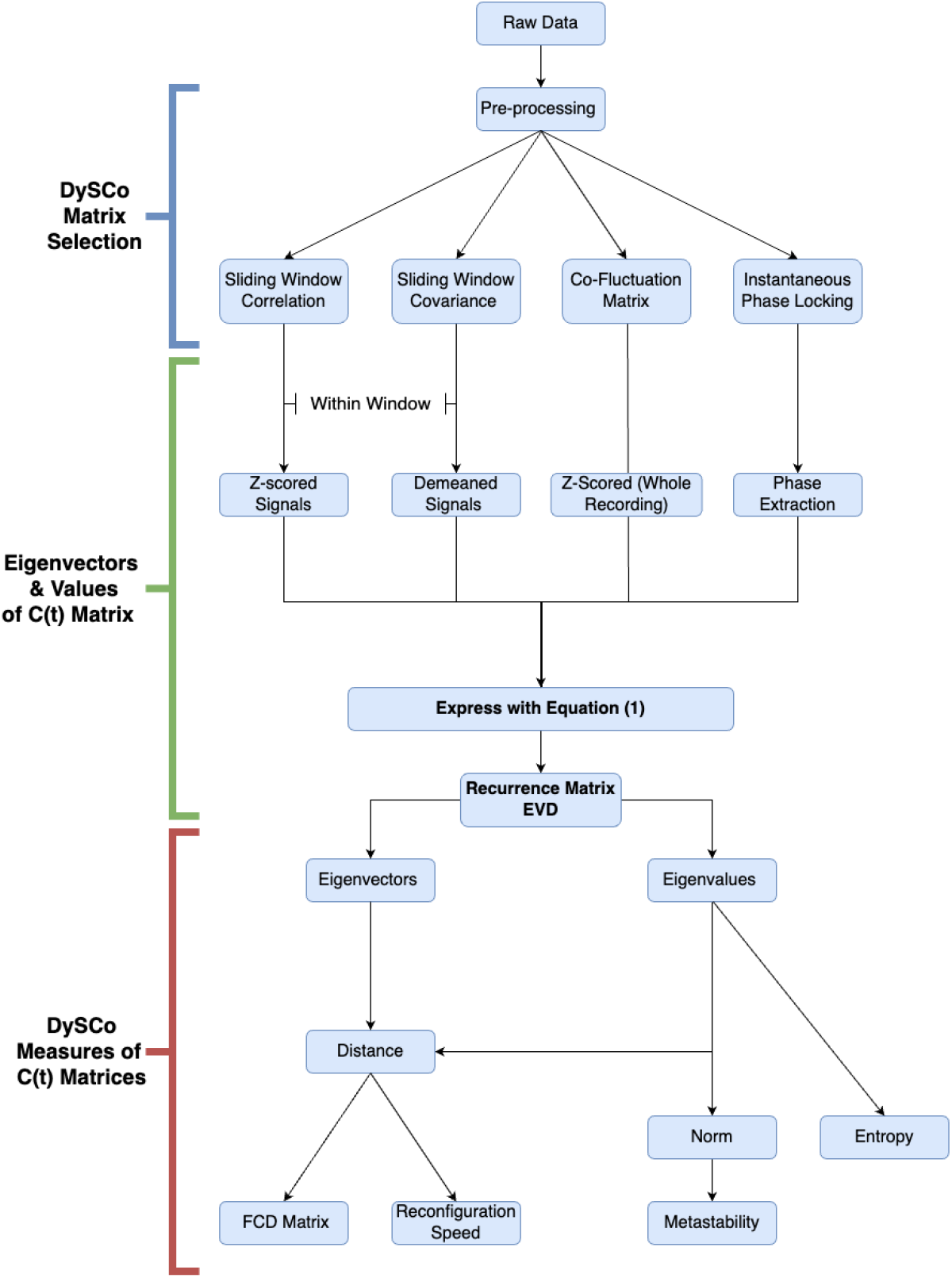
Shows the main steps involved in the DySCo framework as well as important methodological decisions that must be made when using the framework. After input of raw data and appropriate pre-processing there are 4 possible dFC matrices as described in Theory. Based upon the choice of dFC matrix, which we define as **C**(*t*), subsequent processing steps are employed (such as window size adjustment or extraction of phase) to express these dFC matrices into a unified equation (Equation 1). We next calculate the eigenvalues and eigen-vectors associated with the dFC matrices using the Recurrence Matrix EVD. The eigenvalue-eigenvector representation contains all the information needed to perform the dFC analyses, and to compute the DySCo measures described in Theory. The three main measures are Norms, Distances, and Entropy (see 3.6). From them we can obtain derived measures: from the norm it is possible to compute metastability (see 3.6.2), from the distance it is possible to compute the FCD matrix and the reconfiguration speed (see 3.6.3).

**Figure 4:**
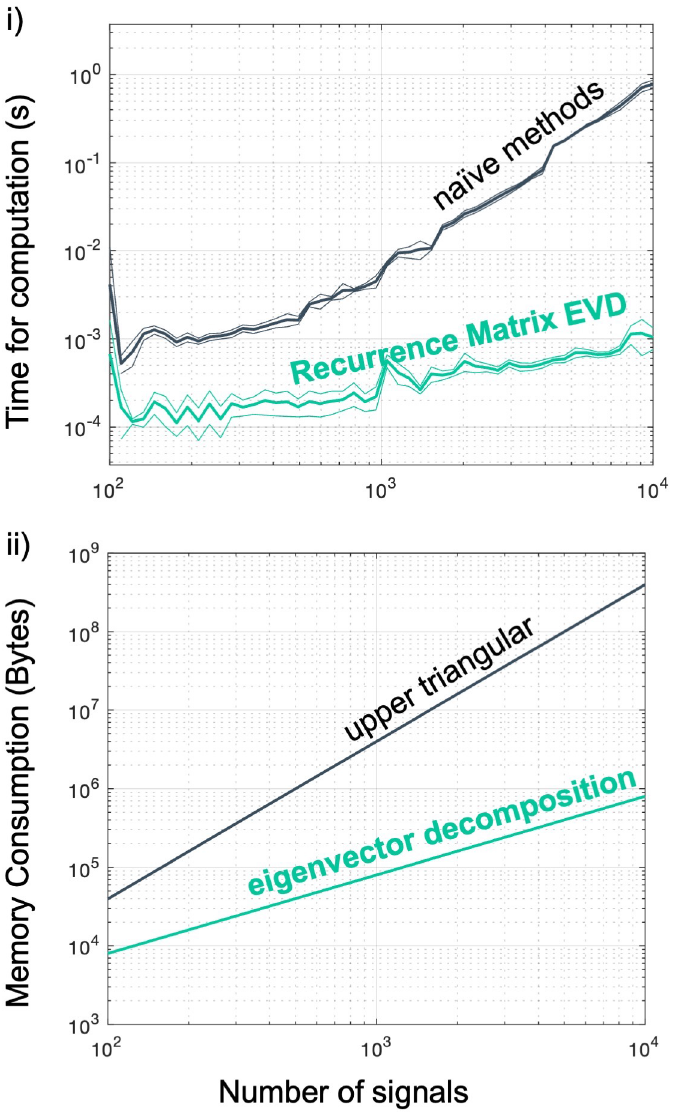
i) Comparison of computational speed of the Recurrence Matrix EVD algorithm compared to generic numerical methods (the MATLAB *eigs* function, see Section 4.2), using randomly generated covariance matrices in a window of size 10. We repeated the experiment 20 times. Thick lines represent the mean computation time, thin lines the *±* variance. ii) Comparison of the memory requirements (in bytes) for the storage of the matrices using their upper triangular form (*N* (*N* − 1)*/*2), versus using the eigenvector decomposition (*NT*).

- the choice of the dFC approach, sliding window correlation/covariance, co-fluctuation matrix or Instantaneous phase locking (see Theory, 1).
- the computation of the eigendecomposition of the dFC matrices using the Recurrence Matrix EVD (see Section 3.5).
- the computation of the DySCo measures based on the eigendecomposition of the dFC matrices (see Section 3.6).

The choice of the type of dFC approach is left to the user and specific use case, as well as the determination of the optimal parameters for the method chosen, such as window size or frequency filtering. In the next subsection, we present an exemplar utilisation of the DySCo framework on the Human Connectome Project.

##### 4.4.2 HCP task fMRI data

We used a sample of pre-processed task-based (working memory N-Back)(fMRI (tfMRI) data from the Human Connectome Project (HCP)(S1200) to illustrate the applicability of this framework to real data. The HCP dataset was chosen as it represents one of the most systematic and comprehensive open-source neuro-imaging datasets available currently. It is a cohesive, well cited and validated, dataset that enables cross-subject comparisons and multi-modal analysis of brain architecture, connectivity, and function and is therefore an effective and suitably diverse dataset upon which to illustrate the DySCo framework.

Detailed overview of the task fMRI acquisition details can be found at ([60]), to summarise: Whole brain EPI acquisitions were acquired with a 32 channel head-coil in a modified 3T Siemens Skyra. Images were acquired with a TR = 720 ms, TE = 33.1 ms, flip angle = 52°, BW = 2290 Hz/Px, in-plane FOV = 208 × 180 mm, 72 slices, 2.0 mm isotropic voxels, with a multi-band acceleration factor of 8 ([61]). During task-based acquisitions two runs were conducted employing opposing phase encoding for each run. Minimal pre-processing is used according to ([62]) but several steps including: removal of spatial distortions, realignment of volumes compensating for subject motion, registration to structural data, bias field reduction, normalization the 4D image to a global mean, and masking of the data with the final brain mask.

In each run n-back task data was collected, which includes 8 blocks of 10 trials per block. Each block commences with a 2.5s cue that describes task type e.g. 2-back or target image (0-back). Within each trial an image is presented to the participant and requires a button press per image. If the image matches the previous image (0-back, 4 blocks of the 8 block design) or was observed two images prior to the current image (2-back, 4 blocks of the 8 block design) then the participant should press a button with their right index finger. If the image is non-matching to either of these conditions then the participant should press a button with their right middle finger. There were a total of 4 distinct categories of images presented (tools, body-parts, neutral faces, and places). Each image category was presented in 2 of the blocks (for both the 0 back and 2-back) across the total 16 blocks in the two runs.

From the HCP pipeline we took each of the different task conditions as described above and generated a single task time course for the HCP Working Memory task (illustrated in Section 5.4) by: (i) convolving each of the time courses for each of the eight conditions with a double gamma canonical hemodynamic response function (HRF); (ii) subsequently, we took the mean value for each TR across all conditions resulting in a single time course.

We randomly sampled 100 pre-processed tfMRI participants from the available 1,034 subjects in the HCP dataset. From these we constructed functional time-series data of the cortical hemispheres using python Nibabel libraries ([63]). As this is simply an illustration of the potential of this framework, we decided to only use the left-hemisphere. This led to a 32492 by 405 matrix corresponding to the total voxels and TRs respectively. We then filtered this array by removing voxels containing either 0 or null values (as these correspond to tissue boundaries).

We expected our measures to temporally align with the task paradigm, confirming their dynamic sensitivity. Specifically, we expected the reconfiguration speed to present peaks corresponding to the switching between task and rest, and the FCD matrix to reflect this task-rest structure. Moreover, we expected the eigenvalue spectrum of the dynamic matrix to be different in task versus rest, which should be highlighted by the Von Neumann Entropy.

#### 4.4.3 Preliminary exploration of the DySCo matrices

The first step is the choice of the most suitable dFC matrix in the DySCo framework to apply to the data. We did that by both visual inspection and quantification of the similarity between different matrices. For the visual inspection, we randomly selected two signals from a subject and plotted the sliding window correlation in time, the sliding window covariance, both with a window size from 5 to 50, the co-fluctuation and the instantaneous phase alignment. See Results (5.3).

We tested two predictions from the Theory: the Instantaneous Phase Alignment Value should be analogous to the sliding window correlation at a specific time-scale of observation (See Section 3.2) and that the co-fluctuation should be analogous to a sliding window covariance with minimal window size (See Section 3.2).

For the equivalence between sliding window correlation and **iPA** at a specific time-scale, we expect that there is a specific window size that maximises the similarity between the instantaneous phase alignment and the sliding window correlation. We expect this peak to be dependent on the bandwidth of the signals. Therefore, we selected 500 random couples of signals for each of the subjects and for each couple of signals we computed the similarity (using Pearson Correlation) between the time trace of the instantaneous phase alignment and the sliding window correlation at all the different window sizes. Note that to be computed the instantaneous Phase Alignment requires the data to be bandpassed. We bandpassed the data in three frequency ranges, 0.01-0.04 Hz, 0.01-0.08 Hz, 0.05-0.1 Hz. We underline that 0.01-0.08 Hz, is the gold standard frequency range adopted for fMRI data ([33, 17, 37]). However, we also tried a lower and higher bandwidth to check our expectation that the bandwidth plays the same role that the window size plays in the correlation matrix.

For equivalence between sliding window of size 1 and co-fluctuation, we repeated the same analysis above to compare co-fluctuation and sliding window covariance with different window sizes. Please note that we did not conduct additional comparisons (for example correlation with covariance) because they represent quantities that are conceptually different.

Based on our investigation (see Results, Section 5.3) we decided to proceed using the sliding window correlation matrix.

#### 4.4.4 Voxel level sliding window correlation analysis of HCP data

Using the results from the preliminary analysis we next applied the DySCo framework to compute sliding window correlation in the whole sample of 100 participants(window size = 21, number of eigenvectors = 10. From the obtained eigenvectors and eigenvalues we calculated the following DySCo measures: Reconfiguration Speed, the Functional Connectivity Dynamics (FCD) matrix, (see 3.6.3), Von Neumann Entropy. To show the sensitivity of the Von Neumann entropy and reconfiguration speed we calculated the correlation between these measures and the original task time-course using Pearson’s correlation coefficient. To assess the sensitivity of these measures at the single-subject level we repeated the analysis on a single example subject chosen from the available participants. In the results we show measures of entropy, speed, and FCD matrices for all subjects as well as at the individual subject. For visualisation purposes we also extracted the 3 eigenvectors associated with the 3 biggest eigenvalues of the dFC matrix.

## 5 Results

In this section, we first show the computational efficiency of the Recurrence Matrix EVD, we then illustrate the DySCo framework on a simulated dataset. We compare the behaviour of the four types of dFC with respect to two exemplar signals, and also confirm theoretical prediction on the equivalence of the sliding window correlation, co-fluctuation and IPA in certain limit cases. Finally, we illustrate the DySCo framework with the analysis of an n-back task dataset from the Human Connectome Project.

### 5.1 Computational Speed and memory consumption

We confirm that the Recurrence Matrix EVD algorithm to compute the eigendecomposition of the dFC matrices offers an extremely significant speed up over conventional methods (see Section 4.2). As shown in Figure 4 i), the RMEVD outperforms naïve numerical methods. It is 100 times faster for matrices with dimensions of 1000×1000, and 1000 times faster for matrices larger than 10,000×10,000. As shown in Figure 4 ii), the matrix decomposition in the *T* eigenvectors also allows to save each matrix with *NT* elements instead of *N* (*N* − 1)*/*2, offering a significant advantage in terms of RAM requirements, without loss of information.

### 5.2 Simulated Dataset

Before turning to real data, we needed to ensure that we could effectively recover the covariance matrices that we underimposed in the simulated data (see Section 4.3). As shown in Figure 5, we were able to correctly recover the ground truth covariance matrices. The Reconfiguration speed shows peaks coinciding with the switches from one matrix to the other. Finally, the Functional Connectivity Dynamics confirms that the timeseries is temporally clustered in the 5 underimposed patterns. Altogether, these results show that the dFC matrices computed with the Recurrence Matrix EVD formulas match the ground truth connectivity patterns and that DySCo’s dynamic measures can quantify their changes in time.

**Figure 5:**
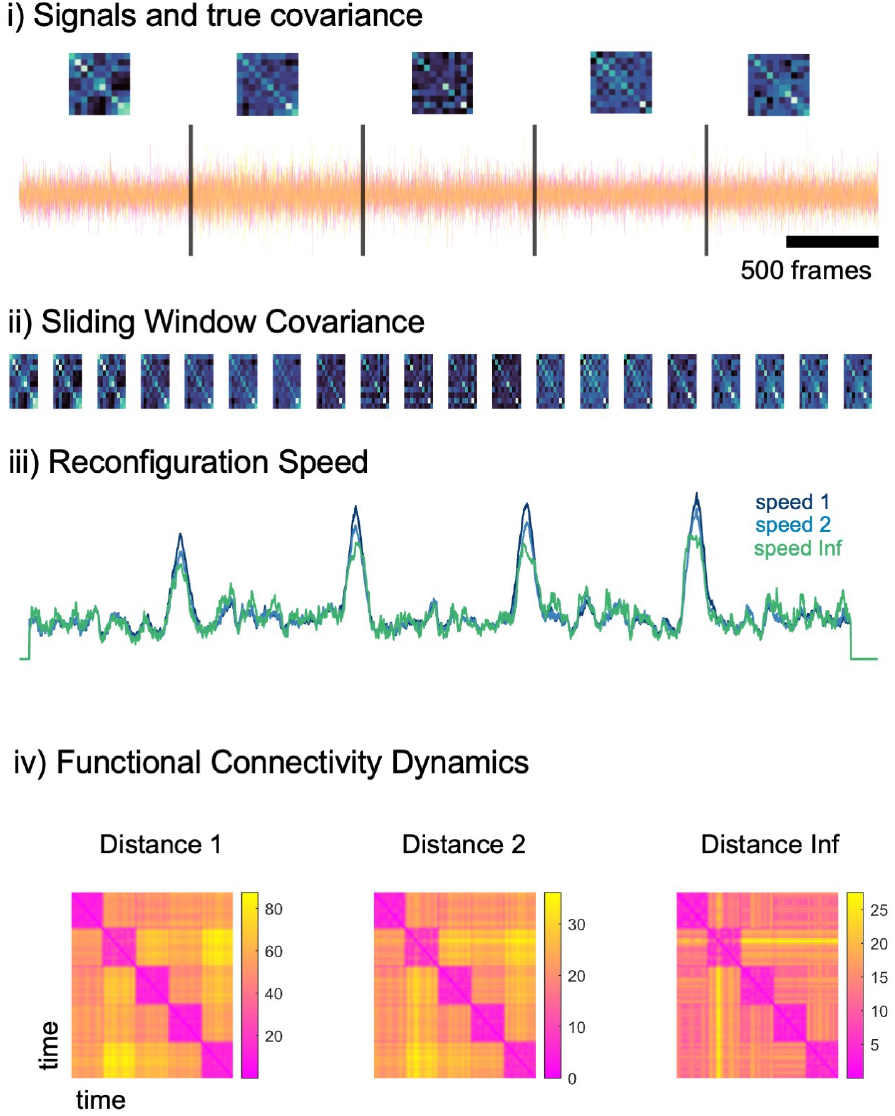
Application of the DySCo measures in a simulated dataset. i) *N* = 10 simulated signals and the five underlying covariance patterns, corresponding to brain states ii) The sliding window covariance matrix computed using the DySCo formula iii) Reconfiguration speed with a lag of 100 frames shows peak corresponding to the switches between brain states iv) Functional Connectivity Dynamics matrices. *FCD*_*ij*_ is the distance between the matrix at time *t*_*i*_ and the matrix at time *t*_*j*_ using the three possible distances proposed in the DySCo framework.

### 5.3 Exploration of the DySCo matrices

#### 5.3.1 Matrix Type

Each type of dFC matrix reacts differently to the same data, and understanding what each is sensitive to is important when making the choice. Looking at Figure 6 A, the sliding window correlation matrix and the covariance matrix convey two different pieces of information, the latter being more sensitive to amplitude changes. As expected, a larger window size, in lighter shades, implies a suppression of high amplitude time-localized correlation/covariance peaks, therefore, losing time sensitivity. As described in the Theory (see Section 3.2), the Instantaneous Phase Alignment value is analogous to a correlation of signals approximated by sinusoid at a specific bandwidth of observation. Indeed, in Figure 6 B.i, we see a peak at a specific timescale where **iPA** is most similar to the sliding window correlation. We also find that this peak depends on the bandwidth of the signal: a lower bandwidth shifts the peak on the right. Indeed, the more low-frequency is the phase alignment, the larger window is needed to capture this with a correlation matrix. Moreover, as quantified in Figure 6 B.ii, the co-fluctuation is similar to a covariance matrix with window size approaching 1.

**Figure 6:**
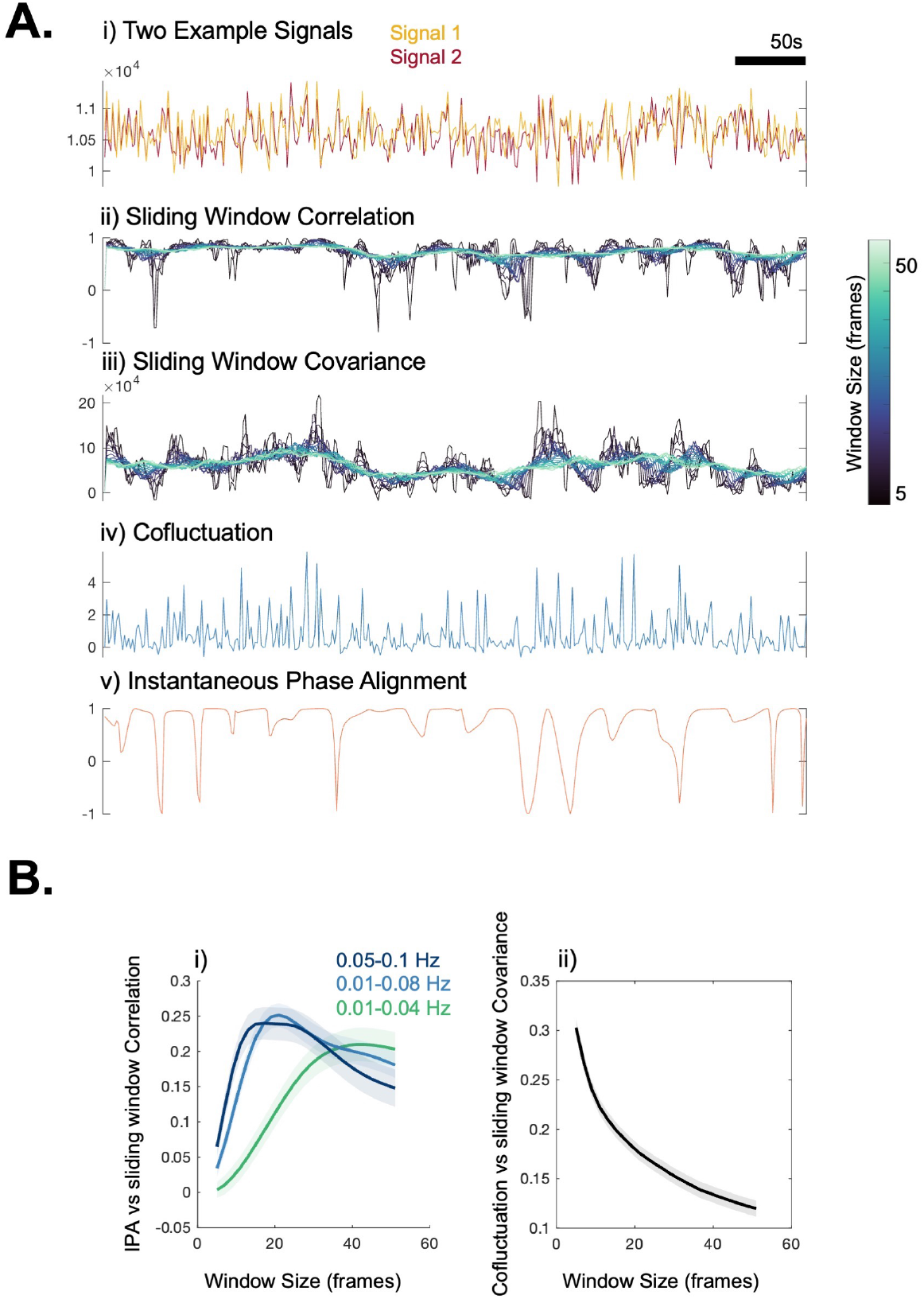
**A**. We selected two random signals from an example participant (i) and plotted the time course of all the measures of the DySCo framework: (ii) sliding window correlation, with window sizes ranging from 5 to 50, (iii) sliding window covariance, with window sizes ranging from 5 to 50, (iv) co-fluctuation, (v) instantaneous phase alignment. **B**. (i) The average (across different couples of signals, across subjects) correlation in time between instantaneous phase alignment and sliding window correlation as a function of the window size. (ii) The average (across different couples of signals, across subjects) correlation in time between co-fluctuation and sliding window covariance as a function of the window size.

This procedure was used as a tool to investigate and choose the matrix for the HCP application. We chose the sliding window correlation matrix based on the following considerations:

1. For the purpose of this work coordinated variation was more relevant than intensity, so we excluded covariance and co-fluctuation.
2. As explained in the Theory, and confirmed by this section, the Instantaneous Phase Alignment matrix assumes a specific, narrow, bandwidth of observation of the signals (timescale), and, moreover, assumes that brain areas are modeled as sinusoids. We did not have justifications for these two hypotheses in our signals, which is why we chose the sliding window correlation matrix.

### 5.4 Application of voxel-level dFC to the HCP dataset

Our final pipeline employed the sliding-window correlation matrix and contained a window size of 21 frames. Matrices were approximated and denoised by retaining their 10 first eigenvectors. As in Figure 7, the reconfiguration speed for all subjects shows the same distinctive peaks as were observed in the simulated data with changes in the task, with the largest peaks shown at the onset of task switching (task/rest) indicating shifts in connectivity patterns between task and rest.

**Figure 7:**
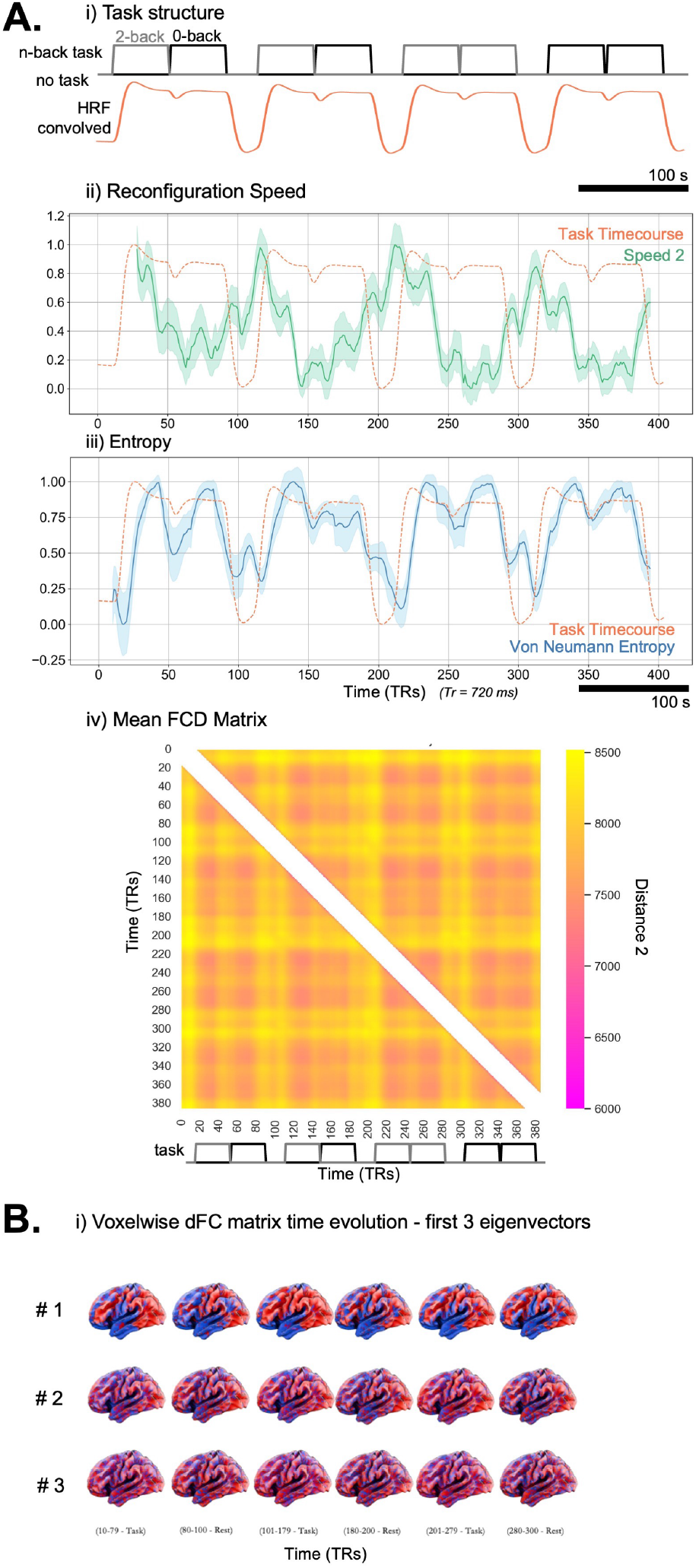
Application of DySCo to HCP dataset (all subjects). **A**. i) the task structure (gray line), and the HRF convolved task timecourse, in orange (see Section 4.4.2). ii) Shows the mean reconfiguration speed (green) standard error (shaded) calculated from the obtained eigenvalues across 100 subjects with a window size of 20. The dashed line again shows the Task time course of the HCP n-back task (r = –0.46, p *<* 0.001). iii) Shows the mean von Neumann Entropy (blue) standard error (shaded) calculated from the obtained eigenvalues across 100 subjects with a window size of 20. The dashed line shows the Task time course of the HCP n-back task (r = 0.76, p *<* 0.001). iv) Shows the FCD matrix averaged across all subjects. The entry *ij* of the FCD matrix (see 3.6.3) represents the distance 2 between the dFC matrix at time *t*_*i*_ and the dFC matrix at time *t*_*j*_. **B**. To show an example of evolution in time of the sliding window correlation matrices, we show them by using their first 3 eigenvectors (averaged across all subjects).

The average Von Neumann entropy showed a strong positive correlation with the task-timecourse (r = 0.76, p *<* 0.001). Also, the averaged FCD matrix across all subjects shows a block structure corresponding to the task timecourse. In Figure 7, we show a way to visualise the extremely high dimensional sliding window correlation matrix (almost 1 Billion entries, impossible to show as a matrix) by means of its 3 leading eigenvectors (plotted on the FSLR projection (32K MSM)). We show how the matrix changes in different moments of the recording.

In Figure 8 we also show the same measures applied to a single, example, subject using the same parameters and matrix choice. The reconfiguration speed here also shows pronounced peaks at the onset of switching from task to rest, but also distinctive peaks during the 2-back task. The Von Neumann entropy measure for a single subject also shows a strong positive correlation with the accompanying task-timecourse (r = 0.89, p *<* 0.001). Also the FCD matrix presents a distinctive block structure corresponding to task time-course.

**Figure 8:**
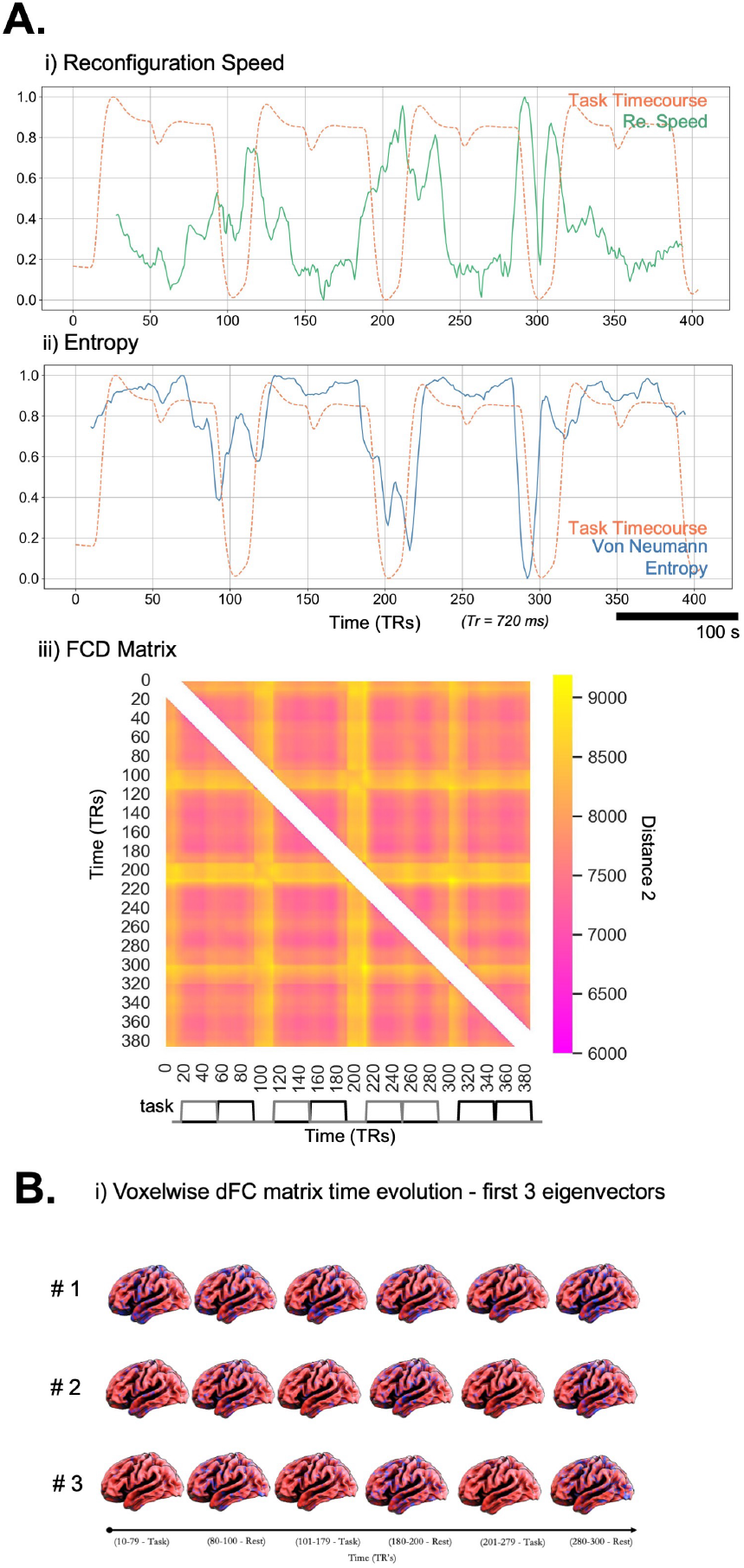
Application of DySCo to HCP dataset (One Example Subject). **A**. i) Shows the reconfiguration speed (green) for the single example subject. The dashed line shows the Task time course of the HCP n-back task (r = –0.66, p *<* 0.001). ii) Shows the von Neumann Entropy (blue). The dashed line shows the Task time course of the HCP n-back task (r = 0.89, p *<* 0.001). iii) Shows the FCD matrix for the single example subject. **B**. To show an example of evolution in time of the sliding window correlation matrices, for the single example subject, we show them by using their first 3 eigenvectors.

We chose the correlation of Von Neumann Entropy with task timecourse as a measure to test the sensitivity of our proposed approach to the change of window size. Supplementary Figure S1 shows the time course of the Von Neumann Entropy using multiple window sizes ranging from 10-24 frames. This shows that our results are robust to a change in window size. The correlation between entropy (at specific window size) and task time-course is also shown and varies between a min of 0.69 (window size = 10) and a max of 0.87 (window size = 19).

## 6 Discussion and Conclusion

We bring the mostly commonly used dynamic functional connectivity (dFC) methodologies into an integrated mathematical framework. This enables the definition of a set of metrics, associated with the eigenvectors and eigenvalues of the dynamic matrices, that have a common underpinning and interpretation. At the core of the framework lies the Recurrence Matrix EigenVector Decomposition (RMEVD) algorithm which allows an extremely fast eigendecomposition of ultra low rank matrices. This is made possible by using only the eigenvectors corresponding to non-trivial eigenvalues, thus retaining all dFC information and allowing the computation of all dFC measures without the need to explicitly reconstruct the matrices themselves ([15]). This opens the way to both the analysis of extremely high dimensional data and real-time applications, which up to now have been considered prohibitive.

Indeed, to illustrate the capabilities of the RMEVD algorithm and the relevance of the metrics we introduce, we present the analysis of an n-back task from the HCP at the voxel level, i.e. with a dFC matrix made of 900M elements, and characterise the spatio-temporal structure of the dFC matrices.

### 6.1 Discussion of the DySCo matrices, the DySCo measures and their related parameters

A unified framework allows for an easy characterisation of the similarities and differences of the dFC methods considered.

Changing the window size for the sliding window correlation and covariance approaches changes the temporal scale of observation ([2]). Indeed, at small window sizes it is possible to see high amplitude peaks, which get suppressed at bigger window sizes. Our results further suggest that the covariance matrix captures the co-fluctuation matrix in the limit of the window size approaching 1, which makes it very sensitive to the co-fluctuation peaks ([29]). The physiological relevance of these peaks and their interpretation depends on the experimental setting, e.g. TR, and is left to experimenter.

As the relation between the instantaneous phase alignment (**iPA**) approach and the sliding window correlation, we found that the instantaneous phase alignment is sensitive to a specific (narrow) time-scale, which is specified by the bandwidth of observation, confirming the observations made in ([33, 17, 37, 15]). Indeed, shifting the frequency of observation shifts the window size at which the maximal similarity with the sliding window correlation was observed. Therefore, when using the **iPA** matrix, the experimenter should be mindful of the bandwidth they are selecting and the implications this has on the signal that is eventually used.

Using the DySCo framework we have introduced 3 important measures of dynamic functional connectivity: i) The norms, which are a measure of the total amount of instantaneous interactions, ii) the distances, which represent the similarity between dFC matrices, and iii) entropy which captures the complexity of the signals. These DySCo measures are extremely sensitive to changes in dFC, as we have demonstrated both in a simulated and real dataset. In summary: the distance captures how dFC evolves in time during the task fMRI recording. The Entropy quantifies the changes in complexity of the signals in the window of observation. Moreover, the measures proposed in the DySCo framework allow to perform all the typical analyses in dFC, i.e., quantification of time-varying total connectivity (e.g. [53]), quantification of the temporal structure of the evolution of dFC matrices ([13]), metastability ([37]), clustering ([4, 17]). This can be done in an unified and computationally optimized manner.

Another advantage of the DySCo framework is that it depends on very few design choices. The most important is the window size. The other choices to be made are related to the norms and distances used, which might be used either for sensitivity analyses, or dependent on experimental settings, task or imaging modality. Another aspect of the pipeline that can be tuned is the number of eigenvectors used to approximate the dFC matrix. The framework as presented here makes a lossless representation possible, but one can chose to focus on a subset of non-trivial eigenvectors. This can be used as a denoising method, if one considers that the information contained in the leading eigenvectors captures the essential signal information.

In this study we chose our input parameters based upon careful exploration of the parameter space, and consistently with the plethora of existing literature into parameter optimisation in these data types ([64]). However, it should be very clear that the parameter sets chosen in this analysis pipeline are specific to the data used and are not directly transferrable to other fMRI datasets or imaging modalities, such as EEG/fMRI/wide-field calcium, and may be dependent on the experimental design.

### 6.2 Example from the HCP data

Using the sliding window correlation matrix, both in individual subjects and across 100 subjects, we have shown that the metrics introduced are sensitive to the changes in the functional connectivity patterns associated with the task.

The reconfiguration speed illustrates the speed at which the matrix **C(t)** evolves with time ([13]). It is therefore capable of capturing temporal representations of connectivity patterns. A distinct spike in reconfiguration speed (Figure 7, 8) reveals a rapid modulation in functional connectivity patterns associated with a switching between task and rest. The reconfiguration speed is able to dynamically track the reconfiguration of connectivity patterns associated to a task not only at the group level, but also at the single individual level. When aggregated across all subjects, a distinct peak is identifiable at the onset of each rest period, with a step-wise increase in peaks occurring across the task period. However, in the single subject we can observe two distinct blocks of increased reconfiguration speed that occur just prior to rest and during the rest period, with a similar, albeit less defined, increase in speed across the task. While it is not suitable to draw inferences as to the meaning of each of these peaks in relation to tangible connectivity patterns, it is clear that this measure is capable of effectively capturing both changes during an activity and between activities. The FCD matrix is also able to summarise across the similarities and differences in connectivity patterns associated with tasks across the whole recording. We observe a distinctive tiled structure that reflects the block task design. During task, all the dFC matrices are more similar distancewise, while rest is associated with periods where the dFC reconfigures itself.

As described in the Theory section, entropy can be viewed as a measure of how broadly the brain explores different axes of variation and is thus related to the dimensionality of the data, as discussed in 3.6.4 (see also [38]). The obtained measures of entropy demonstrate that the eigenvalue spectrum is informative about the dFC matrix structure and that it changes during task. The observed increase in entropy during task represents a dispersion of the eigenvalue spectrum, which indicates increased dimensionality. This may be explained by the fact that during rest the brain is characterised by a dominance of the default mode network (DMN)([65]), which causes the eigenvalue spectrum to be more peaked.

### 6.3 DySCo as a new view on brain dynamics

We believe that the extremely fast computing capacity offered by the RMEVD taken together with the DySCo measures (3.6) and the considerations made in the Theory Section 3.3 pave the way for using dFC as a new way to examine brain dynamics.

A reasonable starting point to study the dynamics of a system in a data driven way is to identify statistical regularities. However, as in argued in [66], the simplest explanation is not necessarily the best. For example, PCA provides a single covariance pattern for a whole dataset, but as pointed out in Figure 1, these axes of variation may change through the time, as the system is not in a stationary state. PCA thus fails to capture dimensions and variations that have a physiological meaning. DySCo provides the capacity to investigate how these axes change in time at scale, instead of defining a unique, static set of dimensions, we enable the study of how the axes themselves change with time. This introduces a new view of dynamic signals, which does not look at the trajectories in a single space, but rather looks at how the space itself changes in time: we directly characterise the dynamic spatiotemporal patterns of interaction, similarly to the approach in ([17]).

We have shown that this view is complementary to the study of how regionwise connections change. The study of dFC requires few assumptions on the signals, and the measures we introduced go at the core general properties of statistical patterns of interaction: their spectral features.

Thus, we believe that DySCo is a timely translational tool that provide highspeed computing capacity that can seemlessly treat the increasing dimensionlity of data generated by new modalities such as from widefield calcium imaging and kilo-scale electrophysiology. This will enable a uniform treatment of dFC across species and fields, given that the core interaction properties of the signals and their evolution in time do not depend on the specific features of the signals, and thus may be preserved across taxonomies (see [24]).

We note also that the DySCo framework and measures can be extended also to other dFC matrices in the future, as long as they are symmetric (for example mutual information matrices, “non-Pearson” statistical correlation…) – such that the spectral theorem holds and the DySCo measures can be computed.

## 7 Acknowledgements

We thank František Váša, Fran Hancock, and Matthew Harvey for the helpful discussions during the development of the framework. We thank Laila Rida and Bryony Goulding Mew for their support. We thank Riccardino Castellano for the insightful discussions about the link between dFC and quantum mechanics.

## 8 Competing Interests

The Authors declare no competing interests.

## 9 Data and Code Availability

The HCP dataset is freely available at www.humanconnectome.org. The DySCO framework comes with a repository. We developed the code to compute all the DySCo quantities both in MATLAB and in Python, which is available at *https://github.com/Mimbero/DySCo*. Note that, both in MATLAB and in Python, we provide all the “core functions” (compute RMEVD, compute norm, distance, etc…) to autonomously build a processing pipeline. However, we also offer an already built example Python pipeline, the one that has been used to process the HCP data. The repository also features a Python GUI to run the analyses.

## 10 Authors Contribution

GdA and AC developed the mathematical framework, under the supervision of PE and FET.

GdA implemented the MATLAB part of the repository, wrote the MATLAB scripts to perform the analyses on the simulated dataset and the preliminary analyses on the HCP signals, with help and supervision by PE.

OS implemented the Python part of the repository, conceived the pipeline to apply DySCO to real data, developed the pipeline to apply DySCo to the HCP dataset, developed the GUI and the functions for graphical rendering, with help and supervision by RL.

OS (for Python) and GdA (for MATLAB) are in charge of the repository and its manteinance.

GdA, OS, AC, RL, JC, FET and PE conceptualised and wrote the paper. PE supervised the overall work.

## 11 Appendix

### 11.1 dFC matrices

We show now in extended form how to compute **C**(*t*) matrices.

#### 11.11. Sliding window covariance and correlation

Let us start from *N* signals *x*_1_(*t*), *x*_2_(*t*), …, *x*_*N*_ (*t*) which compose a vector that evolves with time **x**(*t*) ∈ ℝ^*N*^, in a window of time *t* = 1…*T*.

The estimator of the covariance matrix *Cov* of each couple of signals (i.e. the *ij* element of the covariance matrix) is:

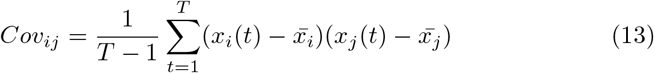

Where 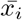 is the mean of the signal *x*_*i*_(*t*) in the window.

If we redefine the signals 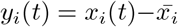, then: 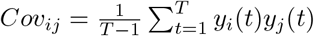.

With a more compact notation, 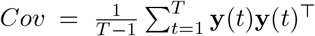, where **y**(*t*) = (*y*_1_(*t*), *y*_2_(*t*), …*y*_*N*_ (*t*))^⊤^.

The same holds true for a sliding window correlation matrix. Here,

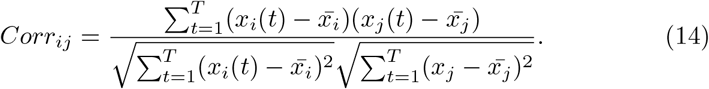

If we z-score the signals, i.e. we redefine the signals *z*_1_(*t*), *z*_2_(*t*)…*z*_*N*_ (*t*), *t* = 1…*T* as the z-score of *x*_1_, *x*_2_…*x*_*N*_, in the window, i.e:

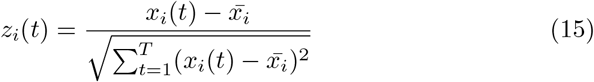

Then:

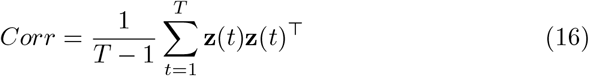

When the sliding window covariance or correlation matrix are computed using a weighted/tapered window, i.e., within the *t* = 1…*T* window of observation the signals are weighted with the positive weights *w*(*t*), *t* = 1…*T*, the equations above hold true, with a preliminary step of weighting the signals **x**(*t*) with the weights *w*(*t*).

#### 11.12 Co-fluctuation matrix

Another widely employed matrix is the co-fluctuation matrix. To compute the co-fluctuation matrix, signals are z-scored to obtain ζ(*t*) in the whole recording (not anymore in the sliding window of size T). Then, the co-fluctuation matrix is:

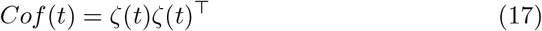

This is analogous to a sliding window covariance matrix where the window size is minimal and equal to 1.

#### 11.13 Approaches based on instantaneous phase differences

When brain areas are modeled as narrow band oscillators, it is possible to define dFC matrices based on the instantaneous phases of signals *θ*_*i*_(*t*). This approach overcomes the limitations of selecting a window of observation of the signals, however, it introduces the need of selecting a band of interest. Indeed, the assumption of narrow band means that a specific time *t*_0_, we can *locally* approximate signals *x*_*i*_(*t*_0_) with oscillators that have the same frequency, an instantaneous amplitude *A*_*i*_(*t*_0_) and an instantaneous phase *θ*_*i*_(*t*_0_). Phases can be extracted using the Hilbert or Wavelet Transform. See ([15]) for an accurate description of this method.

It is of note that the correlation coefficient of two pure sinusoidal waves over their common period is equal to the cosine of their phase shift, i.e.:

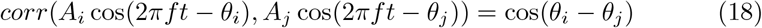

Therefore, once the instantaneous phases of the signals *θ*_*i*_(*t*) are computed, the **iPA** is analogously defined:

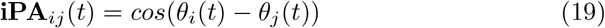

Each entry, similarly to the case of sliding window correlation matrix, is 1 if the signals have exactly the same instantaneous phase, –1 if they are in antiphase, 0 if the two phases are orthogonal. Similarly to the correlation matrix, the **iPA** matrix has a fixed trace of *N*.

Given that the matrix **iPA**_*ij*_ = cos(*θ*_*i*_ −*θ*_*j*_) = cos(*θ*_*i*_) cos(*θ*_*j*_)+sin(*θ*_*i*_) sin(*θ*_*j*_), the **iPA** matrix can be decomposed in a dyadic sum of two terms. We define the “cosine” vector **c** = (cos(*θ*_1_), cos(*θ*_2_), …, cos(*θ*_*N*_))^⊤^ ∈ R^*N*^, and the “sine” vector **s** = (sin(*θ*_1_), sin(*θ*_2_), …, sin(*θ*_*N*_))^⊤^ ∈ ℝ ^*N*^, and rewrite the matrix as:

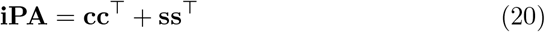

#### 11.2 Eigendecomposition of a dyadic sum

##### 11.2.1 SVD of the data matrix and Recurrence Matrix EVD

Here we relate the eigenvalue decomposition of a dyadic sum 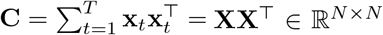 to the singular value decomposition (SVD) of the data matrix **X** ∈ ℝ^*N*×*T*^, and consequently to the eigenvalue decomposition of the recurrence matrix **R** = **X**^⊤^**X** ∈ ℝ^*T* ×*T*^.

The data matrix can be expressed as its SVD decomposition **X** = **UΣV**^⊤^, where **U** and **V** are respectively *N* × *N* and *T* × *T* orthogonal matrices, and **Σ** is an *N* × *T* rectangular diagonal matrix with non-negative entries, the singular values, *σ*_*ii*_ = **Σ**_*ii*_ ([67]). The columns *u*_*i*_ of **U** and the columns *v*_*i*_ of **V** are called left-singular vectors and right-singular vectors of **X**, respectively.

From **C** = **XX**^⊤^ = **UΣΣ**^⊤^**U**^⊤^ and **R** = **X**^⊤^**X** = **VΣ**^⊤^**ΣV**^⊤^, it is straight-forward to observe that:

- The matrices **C** or **R** share the same nonzero eigenvalues, the nonzero diagonal entries of **Σ**^⊤^**Σ** or, equivalently, **ΣΣ**^⊤^, which are the squared singular values of **X**.
- The eigenvectors of the connectivity matrix, columns of **U**, and the eigenvectors of the recurrence matrix, columns of **V**, are the left-singular vectors and the right-singular vectors of **X**, respectively. In particular, the following relation holds **UΣ** = **XV**.

##### 11.2.2 General weighted sum

Here we provide a general proof of the formula for the eigenvector decomposition of a matrix **C** expressed as a weighted dyadic sum:

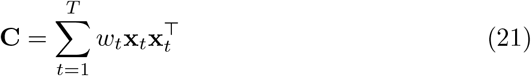

This formula considers both positive and negative weights *w*_*t*_.

This matrix is a linear operator that outputs any input vector into the space spanned by the *T* vectors **x**_*t*_. Therefore, it has rank no greater than *T*. Its first *T* eigenvectors **u**_*i*_ belong as well to the space spanned by the **x**_*t*_ vectors. Therefore, they will be a linear combination of the **x**_*t*_ vectors: for an eigenvector **u**_*i*_, it must be that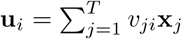. Therefore, the problem of finding the eigenvectors of **C** is equivalent to finding *v*_*ji*_.

By imposing the eigenvector-eigenvalue equation, i.e. **Cu**_*i*_ = *λ*_*i*_**u**_*i*_, and imposing that **u**_*i*_ is a linear combination of the **x**_*t*_, we can write:

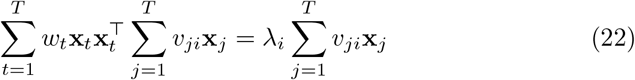

By rearranging terms we can write:

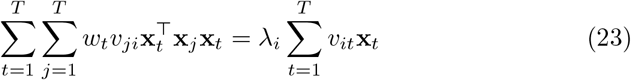

By imposing that the **x**_*t*_ are linearly independent, we have that:

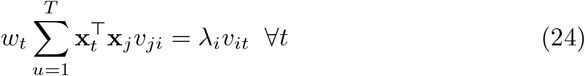

If we define the vector **v**_*i*_ = (*v*_1*i*_, *v*_2*i*_, …*v*_*Ti*_)^⊤^, and the matrix 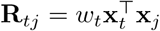equation 24 can be rewritten in matrix notation as:

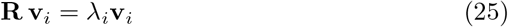

This is an eigenequation for the recurrence matrix. The eigenvalues of the recurrence matrix **R** will therefore be the eigenvalues of the connectivity matrix **C**. The eigenvectors of the connectivity matrix **u**_*i*_ will be related to the eigenvectors of the recurrence matrix **v**_*i*_ by the formula 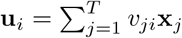

#### 11.3 Computation of distance of dFC matrices in the DySCo framework

##### 11.3.1 General case

Finally, we show here that the distances in the DySCo framework can all be computed from the eigenvector representation of the dFC matrices, without the need of rebuilding the matrices and explicitly computing the norm of their difference.

Having the dyadic representation 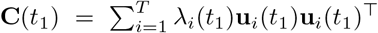 and 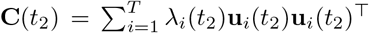 available, we can combine them to express the difference matrix 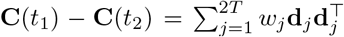, where *w*_*j*_ = *λ*_*i*_(*t*_1_) and **d**_*j*_ = **u**_*j*_(*t*_1_) for *j* ≤ *T* and *w*_*j*_ = −*λ*_*j*−*T*_ (*t*_2_) and **d**_*j*_ = **u**_*j*−*T*_ (*t*_2_) for *j > T*. Therefore, it is possible to resort to the Recurrence Matrix EVD in 11.2 to compute the eigenvalues of **C**(*t*_1_) − **C**(*t*_2_), using a recurrence matrix of size 2*T*. Once the eigenvalues are computed, it is possible to compute the norm of **C**(*t*_1_) − **C**(*t*_2_), i.e. the distance.

##### 11.3.2 Frobenius case

In case of the distance 2, it is possible to compute the distance in an even faster fashion which does not require the eigendecomposition of the recurrence matrix. Indeed, being **C**(*t*_1_) − **C**(*t*_2_) symmetric, it holds:

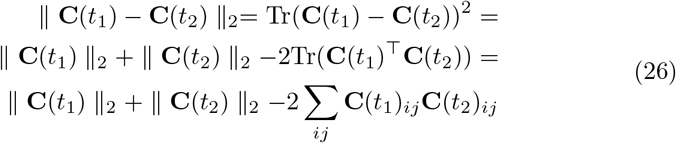

The separated norms of **C**(*t*_1_) and **C**(*t*_2_) are already provided by knowledge of their eigenvalues. We note also that Tr(**C**(*t*_1_)^⊤^**C**(*t*_2_)) is the Frobenius product between the two matrices, which may be a quantity of interest to compute on itself as explained in the main text (Section 3.6.3).

The fast way to compute the Frobenius product, which does not require to expand the matrices, is the following:

Tr(**C**(*t*_1_)^⊤^**C**(*t*_2_)) = Tr(*V* (*t*_1_)*D*(*t*_1_)*V* (*t*_1_)^⊤^*V* (*t*_2_)*D*(*t*_2_)*V* (*t*_2_)^⊤^) by eigenvector decomposition = Tr(*V* (*t*_1_)*D*(*t*_1_)^1*/*2^*D*(*t*_1_)^1*/*2^*V* (*t*_1_)^⊤^*V* (*t*_2_)*D*(*t*_2_)^1*/*2^*D*(*t*_2_)^1*/*2^*V* (*t*_2_)^⊤^) = Tr(*W* (*t*_1_)*W* (*t*_1_)^⊤^*W* (*t*_2_)*W* (*t*_2_)^⊤^) by redefinition of *W* (*t*) = *V* (*t*)*D*(*t*)^1*/*2^. For the circular property of the trace Tr(*W* (*t*_1_)*W* (*t*_1_)^⊤^*W* (*t*_2_)*W* (*t*_2_)^⊤^) = Tr(*W* (*t*_1_)^⊤^*W* (*t*_2_)*W* (*t*_2_)^⊤^*W* (*t*_1_)) =∥ *W* (*t*_1_)^⊤^*W* (*t*_2_) ∥_2_, where *W* (*t*_1_)^⊤^*W* (*t*_2_) is a *T* × *T* small matrix and its 2-norm can be computed by just summing all its squared elements.

#### 11.4 Alignment

Finally, we show that the Frobenius distance between the projector matrices of **C**(*t*_1_) and **C**(*t*_2_) on a subset of their eigenvectors is a measure of alignment of these eigenvectors. Let’s define *U*_*n*_(*t*) as the sub-matrix of the first *n* columns of *U*. The projector matrix on these *n* eigenvectors of **C**(*t*) is *U*_*n*_(*t*)*U*_*n*_(*t*)^⊤^. Thus, the Frobenius distance between the projector matrices of **C**(*t*_1_) and **C**(*t*_2_) is:

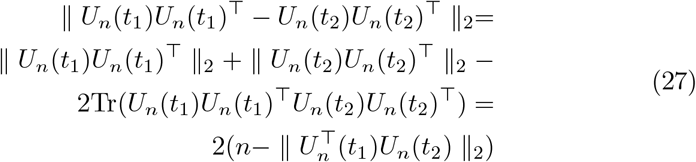

This means that this distance depends on the 2-norm of 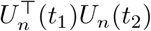, which is a matrix of all the scalar products between the eigenvectors of **C**(*t*_1_) and **C**(*t*_2_). In case the eigenvectors perfectly match, the 2-norm of 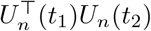 is maximal and equal to *n*, which makes the distance goes to zero. On the opposite side, when eigenvectors are all completely orthogonal the norm of 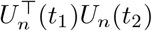 goes to zero making the distance maximal and equal to 2*n*.

#### 11.5 On the estimation of covariance/correlation matrices

A formal problem in dFC is the estimation of high dimensional matrices from a few data points. This problem is inherited from FC analysis that typically suffers from the same limitation, for example when considering low time resolution data such as fMRI: with a parcellation of *N* = 100 regions, there are 2500 entries to be estimated from a few time points, typically a few hundreds.

One explanation why FC can still provide valuable insights is the low enough effective dimensionality of the data: only a small number of eigenvalues are significantly different from 0. This allows to estimate the covariance of a large number of channels using a small number of time points.

DySCo shows that, provided we are considering measures of similarity, it is always possible to see dFC matrices as an *XX*^⊤^ matrix, where *X* is the data matrix. Thus, the eigenvectors are the right singular vectors of the *X* matrix, where *X* has been preprocessed based on what measure is being computed, e.g. for correlation signals are z-scored, for phase locking we are taking the cosine of the angle, etc ….

Thus, doing an SVD of *X* is always formally permitted, even when the number of samples is small. The SVD will still be measuring the geometry of the signals, i.e. the shape of the cloud in Fig. 1, and the DySCo measures quantify how this cloud changes in time. When the window size converges to 1, i.e. in the case of the co-fluctuation matrix, the eigenvector converges to the signal itself. We make it clear that in this case we are studying the sample statistic, which may end up being the population statistic or not, based on the specific application.

The question remains of course whether any real information is captured by the geometry or if it is only noise. Correlation between experimental designs and metrics derived from the DySCo analysis, or any other kind of analysis, is a first validation step. In this paper, we show that indeed not all data is noise and that there is value in the metrics derived from the DySCo framework.

#### 11.6 Sensitivity Analysis

We show in Figure S1 the Von Neumann Entropy with different window sizes.

**Figure S1:**
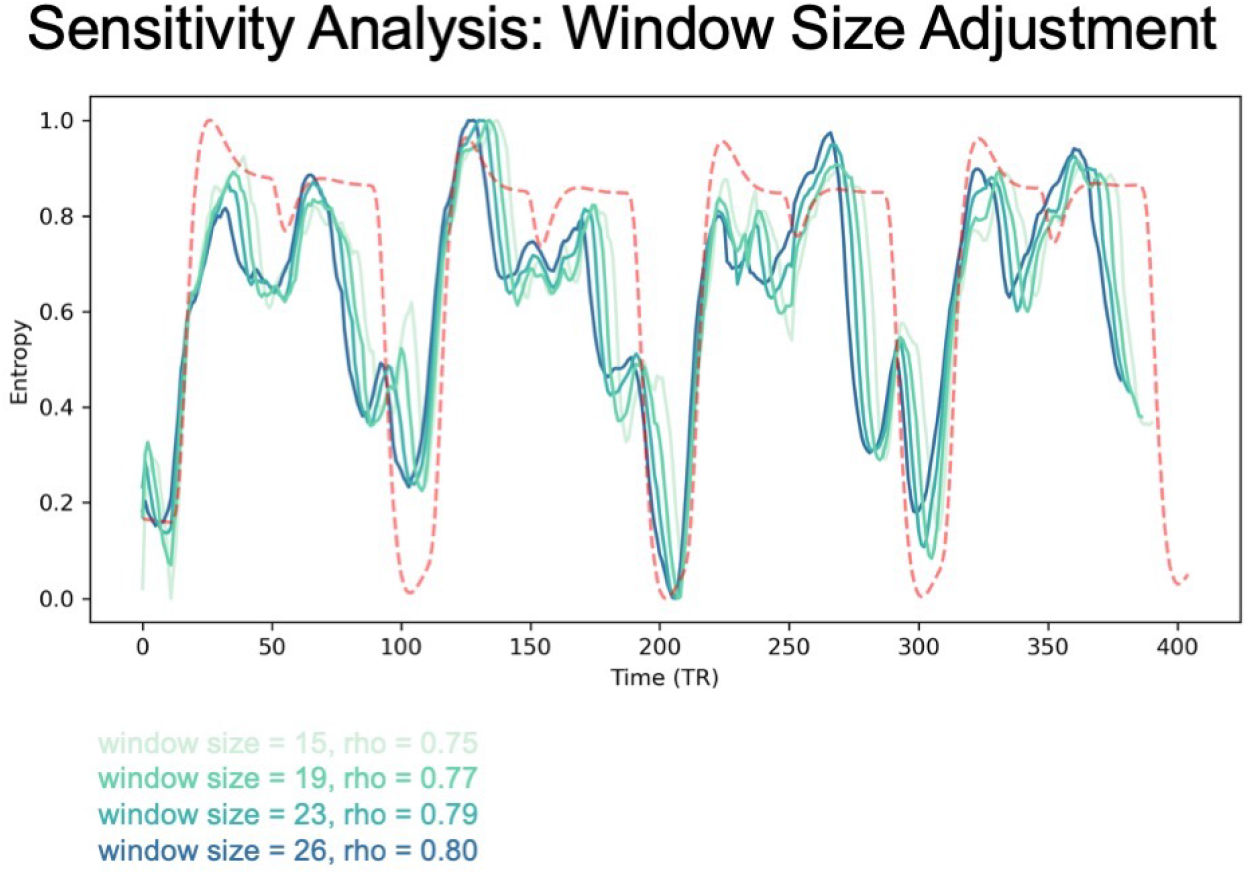
Sensitivity analysis. i) Shows the mean von Neumann Entropy, with each colour of the line denoting the specific window size for the calculation.

## Notes

### Competing Interest Statement

The authors have declared no competing interest.

### Summary of Updates

minor changes to text (intro, discussion, appendix) figure 6 and 7 updated

